# Model-based inference of RNA velocity modules improves cell fate prediction

**DOI:** 10.1101/2023.08.03.551650

**Authors:** Alexander Aivazidis, Fani Memi, Vitalii Kleshchevnikov, Brian Clarke, Oliver Stegle, Omer Ali Bayraktar

**Affiliations:** Wellcome Sanger Institute, Cambridge, CB10 1SA, UK; Division of Computational Genomics and Systems Genetics, German Cancer Research Center (DKFZ), Heidelberg 69120, Germany; European Molecular Biology Laboratory, Genome Biology Unit, Heidelberg 69117, Germany

## Abstract

RNA velocity is a powerful paradigm that exploits the temporal information contained in spliced and unspliced RNA counts to infer transcriptional dynamics. Existing velocity models either rely on coarse biophysical simplifications or require extensive numerical approximations to solve the underlying differential equations. This results in loss of accuracy in challenging settings, such as complex or weak transcription rate changes across cellular trajectories. Here, we present cell2fate, a formulation of RNA velocity based on a *linearization* of the velocity ODE, which allows solving a biophysically accurate model in a fully Bayesian fashion. As a result, cell2fate decomposes the RNA velocity solutions into *modules*, which provides a new biophysical connection between RNA velocity and statistical dimensionality reduction. We comprehensively benchmark cell2fate in real-world settings, demonstrating enhanced interpretability and increased power to reconstruct complex dynamics and weak dynamical signals in rare and mature cell types. Finally, we apply cell2fate to a newly generated dataset from the developing human brain, where we spatially map RNA velocity modules onto the tissue architecture, thereby connecting the spatial organisation of tissues with temporal dynamics of transcription.

## Introduction

The concept of “RNA velocity”, which involves inferring transcriptional dynamics from spliced and unspliced counts in single-cell RNA sequencing (scRNA-seq), has displayed significant potential^1–3^. The first implementations of RNA velocity models^1–3^ have undergone an evolution of conceptual and technical refinements, including improved parameter inference^4–6^, as well as the use of numerical approaches^5,7^,^8,9^ to allow for solving the underlying differential equations. However, these existing refinements are bound to trade-offs between either introducing coarse biophysical approximations^1–4,6,10,11^ or relying on extensive numerical approximations^5,7–9^. Hence, the fundamental challenge remains to define a mathematically sound framework that allows for capturing unconstrained transcriptional dynamics while retaining computational and numerical tractability.

To address the aforementioned limitations, we present cell2fate - a fully Bayesian model of RNA velocity based on a realistic biophysical model of complex transcription dynamics. Cell2fate employs a linearization to decompose complex differential equations into tractable components that can be solved analytically. By doing so, the model is at the same time expressive, interpretable and computationally efficient. The approach to decompose the velocity problem into components also provides a new connection between RNA velocity and dimensionality reduction using a biophysical solution.

We assess and benchmark cell2fate in the context of real-world settings, demonstrating its ability to capture complex dynamics and weak dynamical signals in rare and mature cell types. Finally, we show how cell2fate can be combined with spatial transcriptomics, thereby connecting transcriptional dynamics to their spatial tissue environment.

## Results

### The cell2fate model

Cell2fate builds on a long-stranding history of computational methods for RNA velocity, which employ a dynamical model to explain variation in spliced and unspliced read counts for individual genes and cells (Fig. 1a). At the core of cell2fate is a reformulation of the RNA velocity problem that gives rise to an analytically tractable solution for cell-specific transcription rates. This is achieved by linearizing the corresponding ODE for each gene into a set of M components with simpler dynamics. The dynamics of each module is defined by a switch on/off time, T_m,on_/T_m,off_ on a cell specific time-scale T_c_, as well as corresponding rates λ_m,on_/λ_m,off_ and a gene loading parameter, A_mg_ (Fig. 1b, top right). The linearized solution allows for estimating modules at the level of transcription rates, RNA velocities and observed counts (Fig. 1c, d).

**Figure 1:**
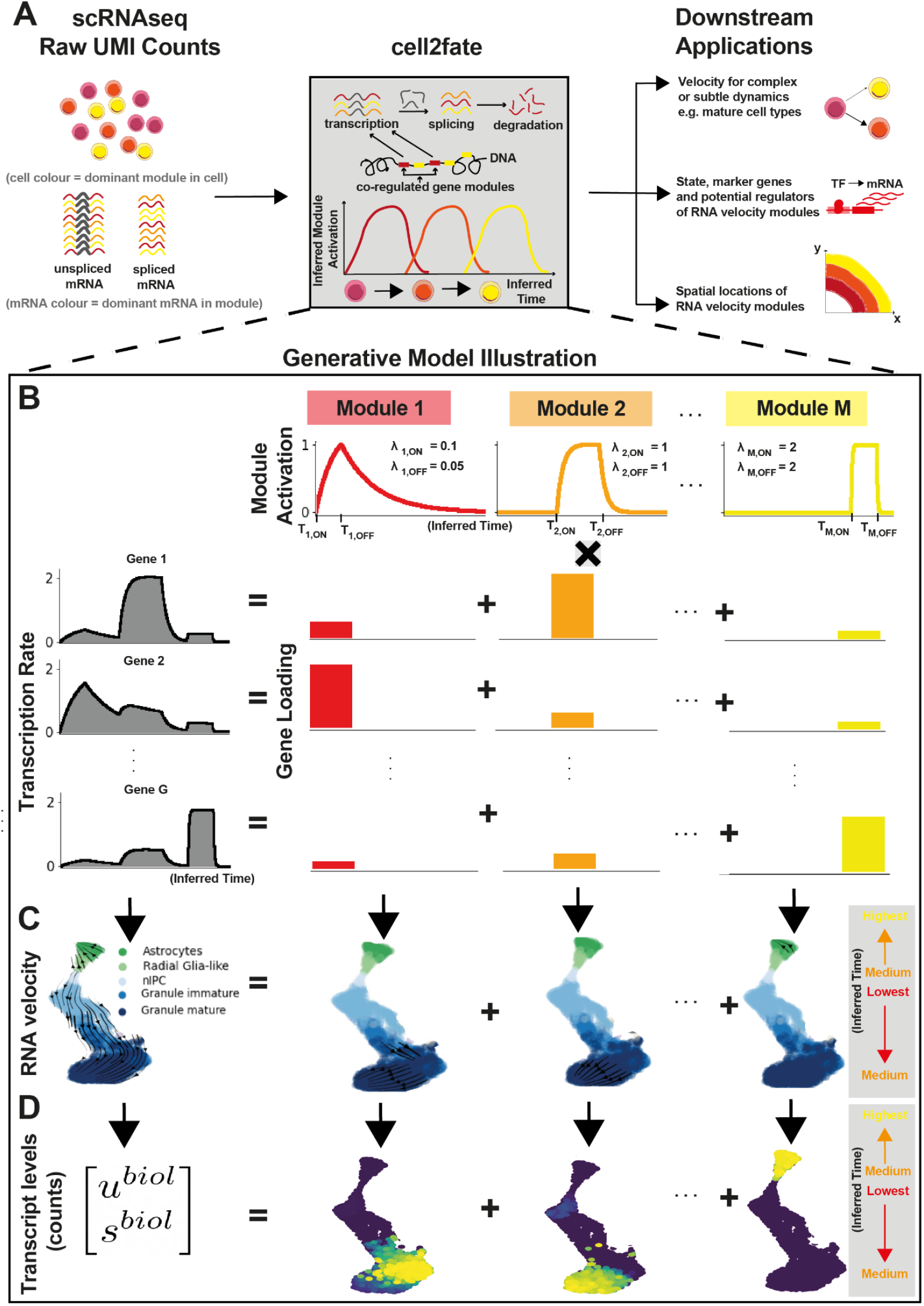
cell2fate model overview. Cell2fate allows to infer complex and subtle transcriptional dynamics by modelling gene-specific transcription rates using a smaller number of independent modules with simple dynamics. **A**, Left: cell2fate input data, comprising raw UMI counts of unspliced and spliced RNA for individual cells and genes; Middle: cell2fate infers time and RNA velocity by modelling gene-specific transcription rates using a small number of modules that explain co-regulated programs; Right: downstream use cases of cell2fate. **B**, Representation of cell2fate as mixed membership generative model. Transcription rates of individual genes (left) are modelled as a weighted combination of prototypical transcription rates of a small number of M modules (right). Each module is defined by a switch on (T_m,on_) and off (T_m,off_) time and corresponding rates (λ_m,on_, λ_m,off_) that determine the speed of activation and deactivation. **C**, Example application of cell2fate to a mouse dentate gyrus dataset covering differentiation of intermediate progenitor cells into neurons and radial glia cells into astrocytes. Left: Overall RNA velocity graph from all modules projected onto a UMAP plot; Arrows denote the total rate of change of RNA across cells. Right: RNA velocity graph of individual modules. Grey box: Time assignment of individual lineages, with the neuron lineages occupying low time points and the astrocyte lineage high time points. **D**, Expected mRNA counts (left) from the cell2fate model, derived as the sum of the analytical solutions of differential equations for each module (right).

The linearization also provides a biophysical connection between RNA velocity and statistical dimensionality reduction. This connection becomes apparent when casting the linearization as a mixed membership model, whereby spliced and unspliced counts of each gene are governed by a linear combination of M modules (Methods, Fig. 1b). The mixing coefficients can then be interpreted analogous to gene loadings of factor analysis or principal component analysis.

The cell2fate model directly operates on the raw cell-level counts as input, and it includes a series of refinements to account for technical sources of variation, including overdispersion, variation in detection sensitivity of spliced and unspliced RNA molecules, ambient RNA and known batches (Methods). Parameter inference is conducted in a fully Bayesian manner, which allows for encoding assumptions on sparsity using hierarchical priors, regularising the effective number of active modules, as well as sharing of evidence strength across genes, cells and modules (Methods). The model is implemented in Pyro and builds on scvi-tools to facilitate its use in existing workflows. The software comes with guidelines and heuristics to determine hyperparameters such as the number of modules (Methods).

### Improved cell fate predictions and estimation of complex of transcription rates

Next, to compare cell2fate to existing RNA velocity methods, we assessed the consistency of estimated cell fate trajectories with prior knowledge. Briefly, we considered the cross-boundary direction correctness (CBDir) metric for benchmarking, thereby scoring the consistency of transition probabilities at the boundary between cell clusters with prior knowledge^3^.

We considered 10 RNA velocity methods spanning different model classes, and approaches for parameter inference (Methods). We applied each method to five scRNA-seq datasets, including widely-used benchmark datasets such as the developing mouse dentate gyrus^12^ and pancreas^13^. In order to assess the ability to resolve complex transcriptional dynamics, we additionally examined mouse erythroid maturation^14^ and human bone marrow^15^, two datasets that feature multiple transcriptional boosts across cellular trajectories^11^. Finally, we considered a mouse bone marrow dataset with markedly low UMI counts^1^, thereby assessing the ability of models to cope with low coverage data.

On average, across all five datasets, cell2fate achieved the best overall performance (Fig. 2a, results on individual datasets shown in Supp. Fig. 1-13). More importantly, cell2fate inferred the correct directionality of cell fate transitions in all datasets, whereas all other methods except for pyroVelocity_model2 inferred a reverse order dynamics in at least one benchmark setting (corresponding to negative CBDir values). Inspecting the benchmarking results, we could attribute the performance of cell2fate to overcoming two major challenges as elaborated below.

**Figure 2:**
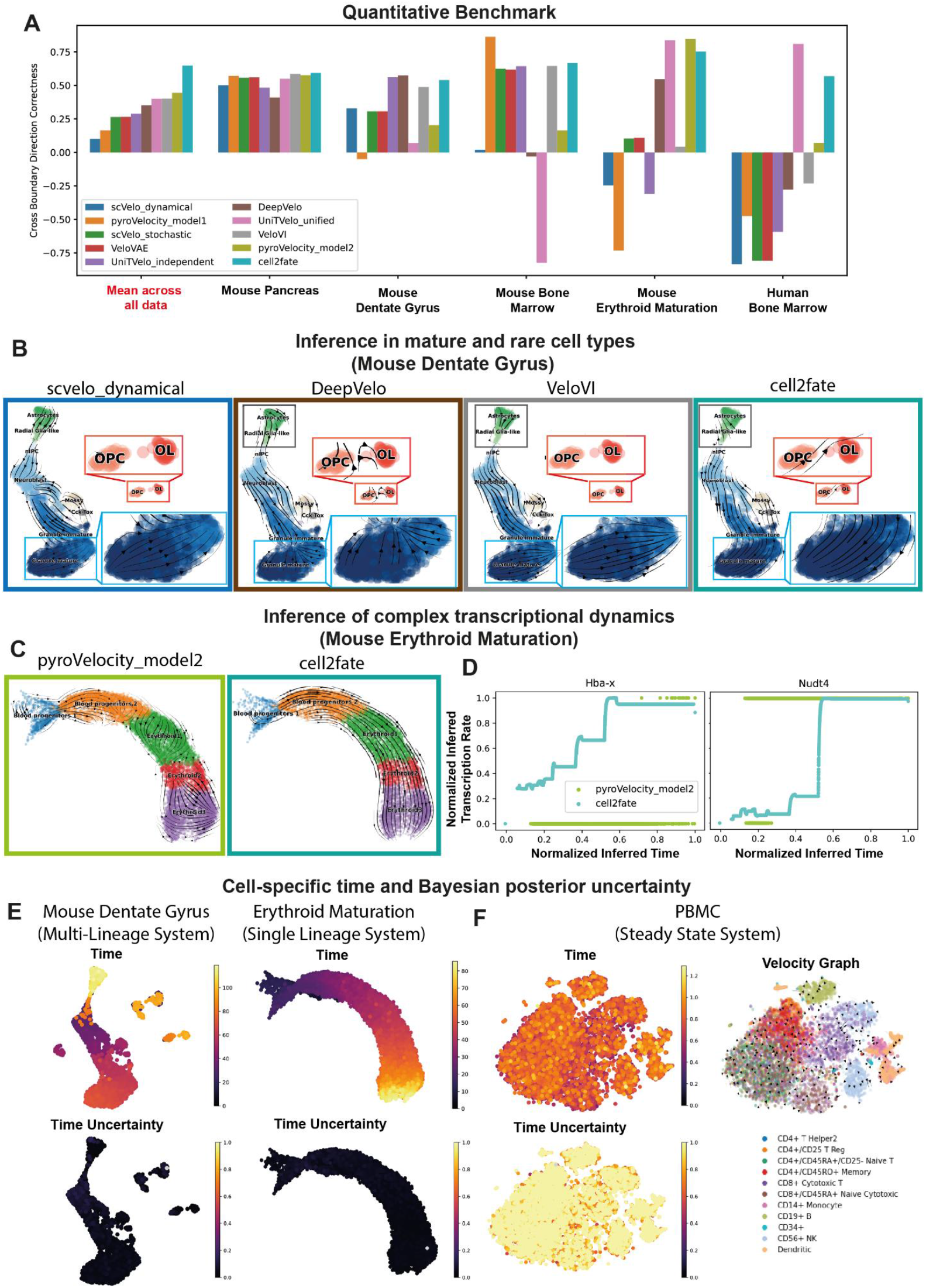
Enhanced performance of cell2fate in RNA Velocity benchmark of 10 methods across 5 datasets. **A**, Performance of 10 methods to reconstruct known trajectories on 5 datasets. Shown is the cross boundary direction correctness^3^ with large positive values corresponding to correct lineage reconstructions; negative values corresponding to opposite directionality. Cell2fate yields the best overall performance across all datasets. **B-D**, Examples of the solutions obtained from selected models and datasets, considering datasets that harbour specific challenges, including (**B**) weak signals in mature or rare cell types and (**C**,**D**) complex transcription rate dynamics. **D**, Transcription rate as inferred by pyroVelocity_model2 and cell2fate for two selected genes that have been postulated to have stepwise changes in transcription rates across differentiation^14^. **E**, Cell-specific time estimates from cell2fate separates different lineages into disconnected time ranges: left, Astrocytes having much higher time than the neuron lineage in the dentate gyrus example; right:, Erythroid Maturation dataset, revealing one connected time range indicating a single lineage. **F**, Coefficient of variation (CV) of the cell2fate cell-specific posterior time can be used as a measure of uncertainty to assess the suitability of a dataset for cell2fate analysis. In a steady state dataset of PBMCs this CV is close to 1 throughout.

First, cell2fate provides sufficient statistical power to identify correct velocity flows from subtle transcriptional dynamics. For example, in the mouse dentate gyrus dataset, other methods consistently failed to resolve the late maturation trajectory of granule neurons and incorrectly suggested that mature cells transitioned into their immature counterparts (Fig. 2b, blue inset boxes, Supp. Fig. 12). Furthermore, alternative methods struggled with rare cell populations such as oligodendrocyte precursors (OPCs) maturing into oligodendrocytes (Fig. 2b, red inset boxes).

Second, cell2fate correctly reconstructs complex transcriptional dynamics. In the mouse erythroid maturation and human bone marrow datasets, the model resolved the correct cell trajectories, whereas other models tended to perform poorly (Fig. 2c, Supp. Fig. 13). Previous analysis^14^, based on the visual inspection of spliced and unspliced counts across manually annotated cell clusters, has provided evidence that mouse erythroid lineage formation features many “multi-rate kinetic” genes such as Hba-x and Nudt4 that display coordinated changes in transcription rates across the cell maturation trajectory^14^. Consistently, cell2fate recapitulated these step-wise transcriptional rate boosts in these multi-rate kinetic genes^14^ (Fig. 2d, turquoise line). In contrast, other methods, such as pyroVelocity_model2 can only predict a single non-zero transcription rate, due to their simpler underlying dynamical model (Fig. 2d, green line). Taken together, these results demonstrate the ability of cell2fate to capture complex cell trajectories and subtle transcriptional dynamics.

We also note that cell2fate provides two additional model outputs that help to provide deeper insights compared to existing RNA velocity methods. First, cell2fate infers a cell-specific time-scale^3,5,6^ (c.f. Fig. 1a,b), which aids the identification of cell lineage progression and distinct cell lineages. For example, in the mouse dentate gyrus dataset, granule neurons and astrocytes are assigned markedly disconnected timepoints, with oligodendrocytes occupying a mid-timepoint range (Fig. 2e, left), consistent with the distinct lineage origins of these three cell types^12^. In contrast, in the mouse erythroid maturation dataset, a single lineage with a single connected time range is identified (Fig. 2e, right).

Second, cell2fate yields Bayesian posterior uncertainty estimates for all parameters, including the time estimate of each cell^4–6^. This provides a principled measure of confidence in the RNA velocity values across and within datasets. In both datasets mentioned above, the coefficient of variation (CV) of the posterior distribution of individual cell times was consistently close to zero, indicating low uncertainty (Fig. 2e, bottom). In contrast, cell2fate applied to a steady state dataset of peripheral blood mononuclear cells (PBMCs)^16^, where no transcriptional dynamics is expected^11^, yields confidence estimates with a CV close to 1, indicating high uncertainty (Fig. 2f). Hence, the CV of posterior cell times can serve as quality control to assess whether cell2fate identifies meaningful dynamics in a given dataset.

Taken together, our benchmark demonstrates cell2fate’s enhanced statistical power to estimate cell trajectories and resolve complex transcriptional dynamics, and the ability to quantify uncertainty in velocity estimates.

### RNA velocity modules reveal fine stages of late cell maturation

cell2fate modules are sequentially activated gene expression programs over time. Given their biophysical foundations in transcriptional kinetics, we expected that RNA velocity modules can provide a more granular characterisation of dynamic processes during cellular differentiation compared to conventional dimensionality reduction techniques that lack a mechanistic basis, such as matrix factorization or clustering. In addition, cell2fate comes with a suite of downstream analysis and visualisation tools, enabling users to explore dynamic processes and derive biological insights.

To demonstrate the cell2fate toolkit, we considered the mouse brain single cell dataset included as part of the benchmarking study (c.f. Fig. 2b), profiling the dentate gyrus region in the hippocampus across two developmental stages^12^. In addition to early differentiation of neurons and astrocytes from neural progenitors, this dataset covers late maturation trajectory of granule neurons (i.e. the late differentiation after the immature neuron stage), a critical process that is however not well understood, and more generally it is unknown whether this late maturation process unfolds across successive transcriptional stages. Previous RNA velocity methods applied to this dataset^2,3^ were able to distinguish neuronal versus astrocyte lineage trajectories, however the correct trajectory for the most mature granule neurons could not be resolved (Fig. 2b).

Cell2fate applied to this dataset revealed 16 distinct RNA velocity modules (Supp. Fig. 2), capturing all the expected cell trajectories, with the dominant lineage corresponding to granule neuron differentiation and maturation stemming from neural intermediate progenitor cells (nIPCs), neuroblasts and immature neurons, while radial glial-like progenitor cells are largely committed to astrocytes (Fig. 3a). We also observed that mossy cells, another neuronal population in the dentate gyrus, were assigned to the middle stages of the granule neuron trajectory. While mossy cells are thought to have different lineage origins, their transcriptional development is highly similar to that of granule neurons^17^.

**Figure 3:**
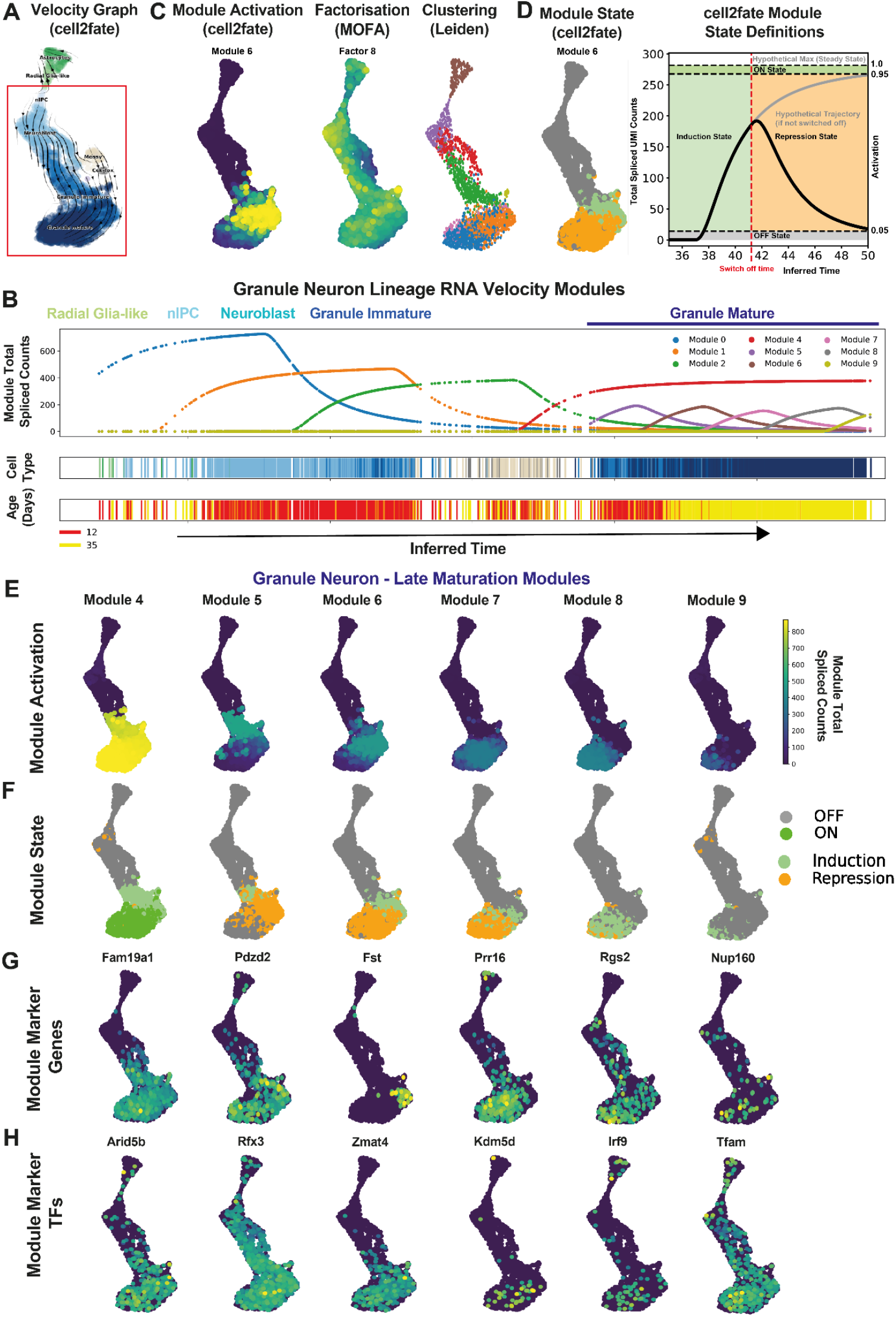
Module decomposition of cell2fate resolves final stages of granule neuron maturation. **A**, cell2fate velocity graph embedding for dentate gyrus data, reproduced from figure 2b for clarity. **B**, Activation of selected example cell2fate module, weight of selected MOFA^18^ (Multi-Omics Factor Analysis) factor and Leiden clustering (with resolution parameter set to 1) for comparison. Overall factor weights and Leiden clusters are more diffusely distributed and less associated with the differentiation trajectory. **C**, Spliced counts abundance caused by selected modules over time. **D**, Module state definitions. < 5% of steady state counts is OFF, >95% is ON. Intermediate values correspond to Induction if T_c < T_mOFF and repression if T_c > T_mOFF. **E**, Activation, defined as the spliced counts produced by a module in each cell. **F**, State of late granule neuron maturation modules. **G**, Module marker genes, defined as having a large part of their transcription rate explained by this module. **H**, Module TF genes, defined as module marker genes that are also known transcription factors.

To explore the dynamics of neuronal differentiation in greater depth, we used the fitted cell2fate model to estimate the total spliced transcript abundance for each of the 9 granule neuron lineage modules in individual cells across the inferred time (Fig. 3b, top panel). This analysis identified the successive induction of modules across the early differentiation of radial glia into nIPCs, neuroblasts and immature neurons (modules #1 to 3). Strikingly, cell2fate also recovered dynamics in mature granule neurons, explained by six modules (modules #4 to 9) that are sequentially activated and temporally overlap across mature granule neurons, thereby finely dissecting the late maturation of these cells into distinct transcriptional windows (Fig. 3b). The model also correctly identified a temporal gap between immature and mature granule neurons (Fig. 3b), which is consistent with prior expectations^12^. The cell2fate visualisation tool complements t-SNE or UMAP by providing dynamic insights anchored on estimated differentiation time, and it can also visualise additional metadata such as cell type annotations or developmental age (Fig. 3b, bottom 2 sidebars).

The total spliced count estimates can also be used to visualise the dynamics of RNA velocity modules across cells, e.g. on a conventional UMAP plot (Fig. 3c). The activation of different modules per cell can be inspected similar to the factor activity in conventional matrix factorization. We also compared these module activation estimates to conventional factor analysis and clustering methods. Briefly, Multi-Omics Factor Analysis (MOFA)^18^ yielded factors that captured complementary sources of variation, with activity profiles that were temporally more diffuse across the differentiation trajectory. Specifically, these factors did not stratify late granule neuron maturation (Fig. 3c, Supp. Fig. 15). We also observed overall low correlation between cell2fate module and MOFA factor gene loadings, particularly for the late neuronal maturation modules 4 to 9 (Supp. Fig. 16). Similarly, Leiden clustering of the scRNA-seq dataset at different resolutions identified clusters that were not aligned with the neuronal maturation (Fig. 3c, Supp. Fig. 17). Collectively, these observations indicate that cell2fate captures complementary aspects of variation compared to existing decomposition methods and is well suited to conduct granular dissection of granule neuron maturation.

Beyond the activity of modules, the dynamics can be further classified into states within each cell, based on whether they are increasing or decreasing in expression (Fig. 3d, Methods). Both quantities are shown in Figure 3e-f for granule neuron differentiation. The additional dynamic information in this visualisation shows for example that module 9 has not reached a steady state, implying that granule neuron maturation continues beyond the time range captured in this dataset.

Finally, we examined to what extent RNA velocity modules can provide deeper insights into the late stages of granule neuron differentiation. We ranked genes by how much of their transcription rate is explained by each module and utilised top genes as “module markers” (Fig. 3g and Supp. Fig. 14). We identified *Rmb24, Fam19a1, Sptb* (Module 4) and *Palm2, Pdzd2, Usp19* (Module 5) as markers switched on in immature granule neurons (Fig. 3g, Supp. Fig. 14). In contrast, *Fst, Rapgef5, Moxd1* (Module 6), *Prr16, Kdm5d, Rpa3* (Module 7) and *Rgs2, Nudt13, 1700048O20Rik* (Module 8) provide novel markers of late granule neuron maturation stages (Fig. 3g, Supp. Fig. 14). *Fam19a1* has been reported to suppress neural stem cell maintenance and promote differentiation^19,20^, consistent with its expression pattern in maturing granule neurons. Rgs2 is dynamically expressed during neuronal activity^21^ and involved in synaptic plasticity^22^, consistent with its late induction in module 8. Apart from these two genes, the marker genes reported here have not been functionally investigated in granule neurons or brain development to our knowledge.

Additionally, we can extract top module marker genes that are transcription factors as “module TFs” (Fig. 3h). Moving on to such module TFs, *Zmat4* (Module 6) and *Tfam* (Module 8) are enriched in late granule neuron maturation stages (Fig. 3h). *Zmat4* has been reported as upregulated in the auditory cortex of young P7 mice compared to adults^23^, while *Tfam* knockouts result in immature neuronal phenotypes^24^. Yet their roles in granule neuron differentiation have not been studied to date. We also find that genes with putative promoter sequences that are most likely to be bound by the top 20 module TFs, as predicted by the ProBound algorithm^25^, are more frequently among the top 300 module genes, than those least likely to be bound by those TFs (Supp. Fig. 18). These TFs provide putative candidate regulators of late granule neuron differentiation.

Taken together, our results demonstrate the great interpretability and statistical power of cell2fate’s module decomposition for single-cell RNA-seq datasets to finely dissect cellular processes and suggest that late granule neuron maturation is composed of distinct stages.

### Spatial mapping of RNA velocity modules

Temporal biological processes are often spatially organised in tissues. For example, cell differentiation and migration are often coupled, with cells associating with distinct spatial signalling microenvironments throughout their differentiation trajectories. Here, we sought to link the temporal information captured by cell2fate to spatial tissue organisation by mapping RNA velocity modules in a newly generated spatial transcriptomics dataset of human brain development (Fig. 4a).

**Figure 4:**
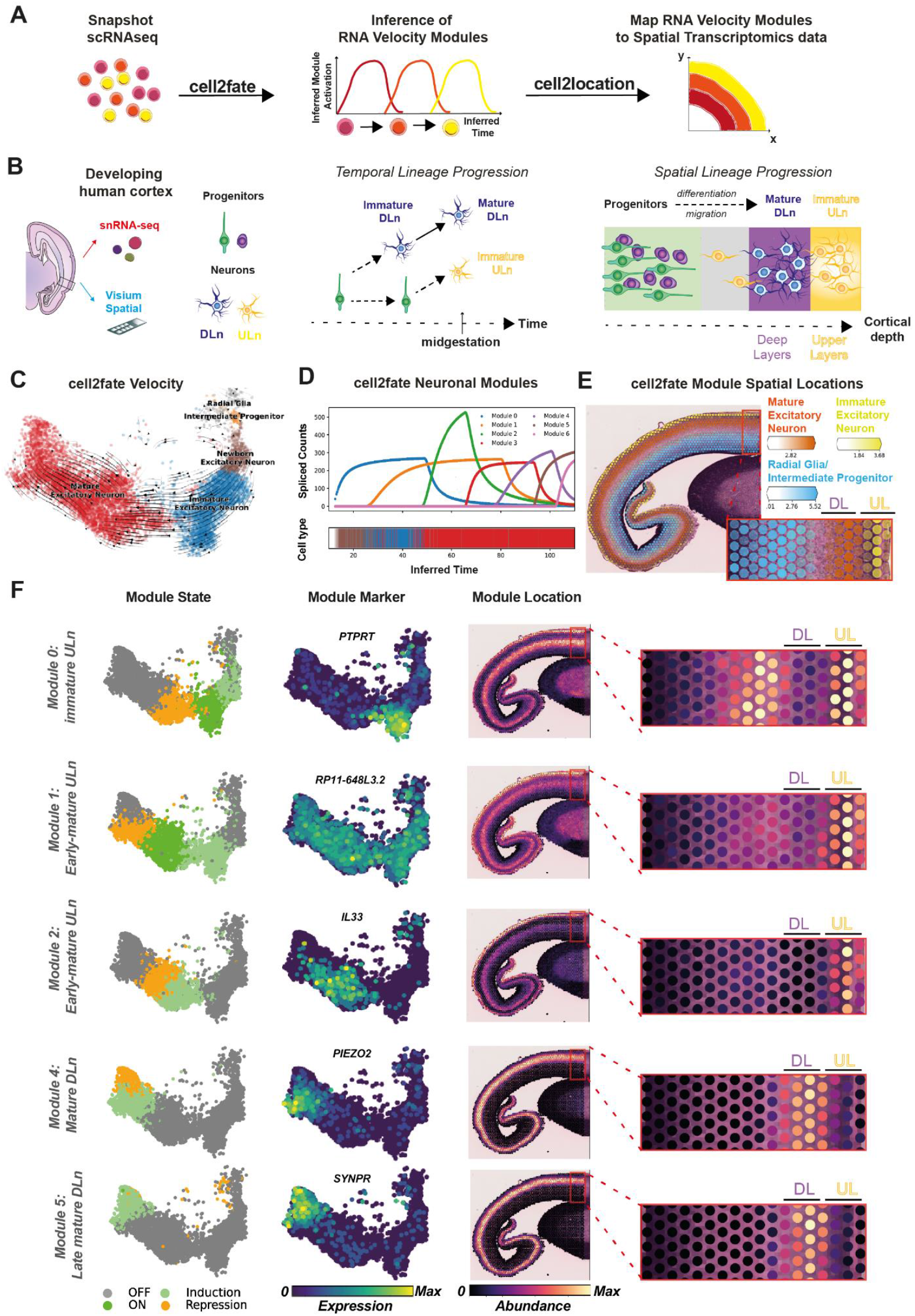
cell2fate interfaces with cell2location to spatially map the cortical neurogenesis process in human brain development. **A**, Workflow for spatial mapping of cell2fate modules. Cell2fate is used to infer modules in scRNAseq data. Steady-state expression of modules is supplied as reference profiles for cell2location to infer abundance of modules across spatial locations. **B**, (left) Illustration of experimental setup: Single-nucleus (10X v3.0) and spatial transcriptomics (10X CytAssist), were performed on cortical tissue sections from two age-matched (13 post-conception week) donors (middle) Illustration of known temporal lineage progression in human cortex: radial glia cells (green) and intermediate progenitors (violet) first give rise to deep layer neurons (blue) early in mid-gestation. They then switch to produce upper layer neurons (yellow) later in mid-gestation, at which point deep layer neurons have matured further. (right) Illustration of known spatial lineage progression in human cortex: progenitors (green, violet) are located deeper inside the cortex, where they produce newborn neurons (yellow), which differentiate and migrate towards the outside, where they form the cortical plate (yellow, blue). **C**, cell2fate velocity graph UMAP embedding for human brain data shows the expected trajectory from radial glia to mature excitatory neurons **D**, Spliced counts produced by each cell2fate module over inferred time illustrate the dynamics of 7 partially overlapping transcriptional programs. Cell type annotations from the UMAP in figure C are visualised by the colour bar at the bottom. **E**, A summary spatial plot of three module locations, named by the cell type in which they reach their steady state expression. **F**, Module state, markers and locations for individual selected modules dissect the spatio-temporal neuron maturation process in detail. Late maturation modules map to deep layers (e.g. modules 4, 5) and early maturation modules to upper layers of the cortex (e.g. module 2).

We focused on the developing human cerebral cortex where excitatory neuron maturation follows a highly stereotyped trajectory through space and time^26^. Neural progenitors termed radial glia and intermediate progenitors reside in the cortical germinal zones, where they sequentially give rise to distinct neuronal subtypes that subsequently migrate out to the deep and upper layers of the cortical plate across their maturation (Fig. 4b). Deep layer residing neurons (DLn) are born before upper layer neurons (ULn) in early gestation, hence DLn are relatively more mature than ULn by mid-gestation (Fig. 4b). Thus, the maturation state and spatial location of cortical excitatory neurons are tightly linked.

To examine cellular differentiation trajectories in the human cortex, we initially performed snRNA-seq profiling (10X v3.0) of one donor at mid-gestation. We then followed standard snRNA-seq processing workflows (Methods) to cluster cells and annotated cell types using markers from literature^27^. We annotated distinct neural progenitors (radial glial and intermediate progenitor cells) as well as excitatory neuron populations at different stages of maturation (Fig. 4c). As expected, mature neurons expressed DLn markers, whereas newborn and immature neurons showed enriched expression of ULn markers (Supp. Fig. 19). We also annotated inhibitory neurons and glial cell types but excluded them from the subsequent excitatory neuron trajectory analysis.

We then applied cell2fate to this human brain snRNA-seq dataset and observed the expected excitatory neuronal differentiation trajectory from neural progenitors to newborn, immature and mature neurons (Fig. 4c). The RNA velocity modules dissected the neuronal trajectory into finer-grained maturation stages, identifying 7 sequentially activated and temporally overlapping modules throughout immature and mature neurons (Fig. 4d, Supp. Fig. 20). While these modules contained some DLn and ULn cell type markers, they also included many genes that are widely expressed across all excitatory neurons in the adult human cortex^28^, such as *PSMC3, KRR1* and *BMPER* (Supp. Fig. 24). This suggests that the modules partially identify a neuronal maturation trajectory common to both DLn and ULn.

In contrast to cell2fate, other RNA velocity methods such as scVelo were not able to accurately identify velocity flow in mature neurons (Supp. Fig. 22). Additionally, the integrated measurement model of cell2fate allowed us to factor in different detection probabilities for spliced and unspliced counts and correct batch effects in our human brain snRNA-seq dataset (Supp. Fig. 21), which is crucial for estimating true transcriptional dynamics from observed counts (Methods).

To spatially map our RNA velocity modules in the developing human cortex, we performed Visium spatial RNA-seq profiling (10X CytAssist) of one cortical tissue section from an age matched donor (Fig. 4b). As the Visium assay offers a coarse spatial resolution and profiles multiple cells at each tissue location (i.e. Visium spot), we used the cell2location algorithm^29^ to deconvolve the abundance of RNA velocity modules across spatial data. We used the steady-state expression counts of each module as reference gene expression signatures, then applied the standard cell2location workflow to infer the abundance of each module signature across Visium spots (Fig. 4a, Methods).

The RNA velocity modules showed expected patterns of spatial mapping across the human cortex (Fig. 4e,f). Progenitor modules spatially mapped to germinal zones (Fig. 4e) while neuronal modules primarily mapped to the cortical plate (Fig, 4e). The fine spatial locations of neuronal modules were consistent with their maturation state (Fig. 4e,f). The immature ULn module (#0) mapped to the upper cortical layers as well as the subplate/intermediate zone that immature neurons pass through during their migration to the cortical plate^30^. The early-mature ULn modules (#1, 2) were exclusively mapped to the upper cortical layers. In contrast, the mature and late mature DLn modules (#4,5) specifically mapped to deep cortical layers.

Taken together, our approach provides a new workflow to spatially resolve complex cell trajectories through tissues.

## Discussion

Here we presented cell2fate, a Bayesian model of RNA velocity that is capable of inferring transcriptional dynamics in settings of complex changes or weak signals in rare and mature cell types. A core innovation of cell2fate is a formulation of the velocity problem that builds on linearization, which allows for solving a biophysically accurate model using analytically tractable linearized components. Another benefit of this formulation is that these linear components can be inspected as interpretable RNA velocity modules. This provides for a direct biophysical connection between cell2fate and statistical dimensionality reduction methods. We illustrated this feature by characterising late maturation trajectories in granule neurons that have been elusive with other methods. Furthermore, RNA velocity modules can be used to locate differentiation trajectories in spatial transcriptomics data. We exemplified this in the developing human brain where the RNA velocity modules of neuronal differentiation showed a high degree of spatial organisation.

The concepts proposed in cell2fate are general and give rise to several extensions that could be considered in the future. It is possible to formulate RNA velocity models with cell-specific splicing and degradation rates, stochastic rates at lineage branching points and causal connections between transcription rates at different time points, equivalent to dynamic gene regulatory networks (Methods). In the long term, dynamical models should also include the effects of cell-cell interactions, based on signalling molecules measured with spatial transcriptomics. An immediate step towards this goal would be combining RNA velocity module mapping with spatial cell-cell interaction tools, such as NCEM^31^, which could identify putative interactions that drive specific steps of a differentiation process.

## Methods

Methods and a description of the cell2fate model can be found in the supplemental methods document.

## Code availability

All results from the cell2fate method can be reproduced with the notebooks included in the cell2fate repository on github: https://github.com/BayraktarLab/cell2fate/tree/main/notebooks

Benchmarking results for all methods can be reproduced with the notebooks in this separate repository: https://github.com/AlexanderAivazidis/fate_benchmarking

## Data availability

Raw UMI counts and metadata in anndata format for all single cell and Visium data is available for download on this portal: https://cell2fate.cog.sanger.ac.uk/browser.html

We will deposit FASTQ files for the human brain single-nucleus and Visium data on ENA by the date of publication.

## Acknowledgements

We gratefully acknowledge Leopold Parts, Yuanhua Huang, Mingze Gao and Chen Qiao for valuable discussions on the cell2fate model, and Elena Prigmore and Jing Eugene Kwa for assistance with human brain single nucleus transcriptomics. This work was funded by the European Commission (ERC project DECODE, 810296) to O.S. and Wellcome Sanger Institute core funding (220540/Z/20/A) to O.A.B..

## Author contributions

A.A. conceived of the cell2fate model, implemented and tested it and produced all figures and results in the manuscript. F.M. generated the human brain single-nucleus and Visium data. V.K. contributed to model conception and implementation on pyro. B.C contributed to the model conception. O.S and O.A.B. co-supervised A.A.. A.A., O.S. and O.A.B. co-wrote the manuscript with feedback from all authors.

## Supplementary Materials

**Supp. Figures 1-25**

**Supp. Table 1: Ground truth transitions and CBDir scores for all methods on Dentate Gyrus Dataset**

**Supp. Table 2: Ground truth transitions and CBDir scores for all methods on Pancreas Dataset**

**Supp. Table 3: Ground truth transitions and CBDir scores for all methods on Mouse Bone Marrow Dataset**

**Supp. Table 4: Ground truth transitions and CBDir scores for all methods on Erythroid Maturation Dataset**

**Supp. Table 5: Ground truth transitions and CBDir scores for all methods on Human Bone Marrow Dataset**

## 1 Cell2fate model description

### 1.1 Background and introduction to the RNA velocity problem

RNA velocity describes the rate of change of RNA in a cell [13], thereby giving rise to a dynamical, i.e. time-dependent models of gene-expression. Popular RNA velocity models to date, such as velocyto [13] and scvelo [1], are fit to spliced and unspliced scRNA-seq counts. The temporal information in the splicing process then allows estimating the direction and rate of expression changes across cells. To achieve this, RNA velocity models formulate a function for the rate of change of mRNA molecules over time. The first methods [13, 1] constructed this function by assuming that each gene produces mRNA molecules with a time-dependent gene-specific rate *α*_*g*_(*t*), unspliced molecules, *u*_*cg*_, are modified into spliced molecules, *s*_*cg*_, with a constant gene-specific rate *β*_*g*_ and degraded with a constant gene-specific rate *γ*_*g*_, as summarized below:

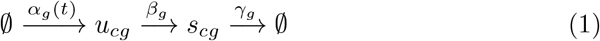

For modelling this process, these differential equations have been suggested [22]:

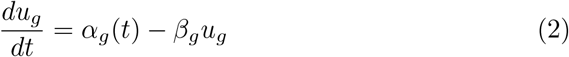

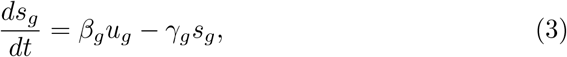

with a step-wise transcription rate that depends on a cell and gene specific time, *t*_*cg*_, and gene-specific switch times *t*_*gON*_ and *t*_*gOFF*_ [13, 1]:

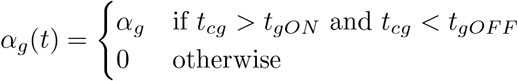

With such a step-wise transcription rate function, the differentiatial equations can be solved analytically [1], which gives rise to time-evolution of spliced and unspliced counts with parameters that can in principle be fit to the observed data. Since parameter fitting is challenging, most methods use various “tricks”, so that models are not directly fit to raw counts. This includes normalizing by total counts in each cell and averaging counts across nearest neighbours in PCA space before parameter fitting. [1].

As noted previously [8, 2], while computationally convenient and tractable, this RNA velocity formulation makes coarse simplifications about the underlying transcriptional dynamics:

1. all rates are deterministic
2. the splicing and degradation rates are constant
3. splicing occurs in a single step
4. the transcription rate can only change to a single non-zero value
5. there is no interaction between transcription at different time points (no gene regulation)
6. genes specific rates are on different time-scales In addition to these conceptual simplifications, the count-preprocessing, implies the following assumptions:
7. total cell-counts are proportional to count detection probabilities
8. there is no ambient RNA in cells
9. there are no batch effects
10. nearest neighbours in PCA space can be treated as nearest neighbours in time

These limitations and assumptions have also been implicated with the variable performance of velocito and scvelo [8, 2] for predicting cell fates.

Following these seminal implementations of RNA velocity, efforts have been directed towards improving the inference procedure [9, 7, 15] or fitting more realistic differential equation models [6], including cell-specific transcription rates [5, 9] and variable splicing and degradation rates [4]. Since the resulting complex systems of differential equation are not analytically tractable, current strategies are based on numerically approximating their solutions, which can impair computational scalibility, accuracy, statistical power and interpretability (see figure 2 in main text and section 3, “Comparative analysis of RNA velocity methods”).

Cell2fate follows a different route and is based on a new formulation of the RNA velocity problem (see section 1.3), which can overcome the limitations listed above without the need for numerical solvers in more complex RNA velocity models. Specifically, we suggest to approximate non-integrable functions, by a sum of integrable components, which we call modules. We use this principle to construct an analytically solveable RNA velocity model with arbitrarily varying transcription rate over time (section 1.3). In addition, we formulate a comprehensive measurement model for spliced and unspliced counts that dispenses with the need for pre-processing counts (section 1.4). This model is then implemented in the probabilistic programming language pyro [3], so that parameters can be obtained via stochastic variational inference (sections 1.5, 1.7). We also give an outlook in the appendix (section 4.5) for how the remaining simplifications can be removed using this linearization approach in the near future.

### 1.2 Definition of cell2fate parameters and subscripts

Symbols for observed data, model parameters and subscripts are defined and explained in detail throughout the text. They are also listed here for additional clarity.

We first define subscripts, which are used to maintain clarity in the dimensions of observed data and model parameters. Capital letters denote the maximum value of the index.

- *c* cells, *C*
- *g* all genes, *G*
- *i* states (ON or OFF), *I*
- *m* modules, *M*
- *j* mRNA maturity (unspliced or spliced), *J*
- *e* experimental batch (usually separate 10X scRNAseq reactions), E

The following symbols denote observed data:

- *U*_*cg*_ unspliced count matrix
- *S*_*cg*_ spliced count matrix
- *X*_*cgj*_ combined count matrix (spliced and unspliced)
- *H*_*ce*_ a one-hot categorical assignment of cells to known experimental batches

The following symbols denote biological model parameters. Time-dependent parameter are either denoted with an additional “(*t*)” or equivalently by adding an additional _*c*_ subscript, since time is a cell-specific parameter.

- *T*_*c*_ the time of each cell
- *u*_*g*_(*t*), *u*_*cg*_ biologically expected unspliced counts
- *s*_*g*_(*t*), *s*_*cg*_ biologically expected spliced counts
- 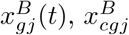 biologically expected unspliced and spliced counts
- *α*_*g*_(*t*), *α*_*cg*_ the transcription rate of a gene
- *β*_*g*_ the splicing rate of a gene
- *γ*_*g*_ the degradation rate of a gene
- *v*_*g*_(*t*) the RNA velocity of a gene
- *α*_*mg*_(*t*) transcription rate of each module
- 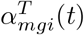 the target transcription rate of each module
- *λ*_*mi*_ rate of transcription rate change for each module
- *T*_*mON*_ switch ON time of each module
- *T*_*mOFF*_ switch OFF time of each module
- *A*_*mgON*_ target transcription rate in the ON state

These are the measurement model parameters:

- *s*_*egj*_ ambient RNA counts
- *l*_*cgj*_ detection efficiency
- 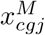 expected counts after accounting for measurement noise
- *a*_*gj*_ Negative Binomial overdispersion parameter

### 1.3 cell2fate biological model definition by linearization

Following previous approaches, we aim to model RNA velocity with more general differential equations, in which the transcription rate is itself going through dynamic changes parameterized here by a function *F*_*α*_(*t*)

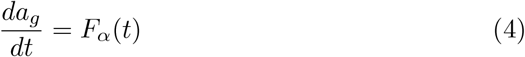

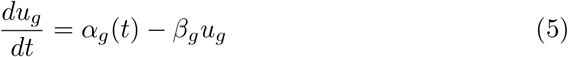

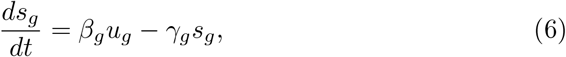

where RNA velocity *ν*(*t*) is the time-derivative of spliced mRNA:

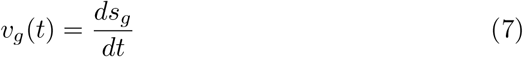

Gene transcription rates are generally believed to be a complex non-linear function of changes in active transcription factor abundance in the nucleus [21], where activation states of transcription factors can be driven by signalling events from neighbouring cells [19]. At the core of cell2fate is the concept to approximate this complexity with a linear sum of easily integrable functions:

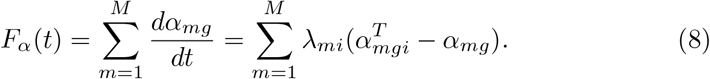

Each component in the sum is denoted by a subscript *m* and is denoted a *module* in the following. The state index *i* can take on the two values *ON* and *OFF* (active or inactive state), depending on a time assigned to each cell, *T*_*c*_, and so-called module switch-times, *T*_*mON*_, *T*_*mOFF*_ :

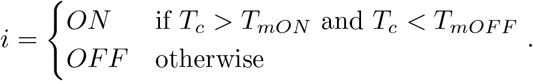

The parameter 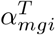, which we call the target transcription rate takes on the following values in each state:

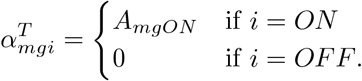

The parameters *T*_*c*_, *T*_*mON*_, *T*_*mOFF*_, *λ*_*mi*_ and *A*_*mgON*_ are determined from the data with prior distributions defined in section 1.5. We physically interpret the transcription rate function by postulating that within small time windows defined by *T*_*mON*_, *T*_*mOFF*_, changes in the transcription rate are driven by changes in only a small number of active regulatory proteins (transcription factors or co-factors) that simultaneously increase or decrease in abundance. The transcription rate function captures the effects of such increases or decreases up to a saturation point defined by 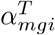 .

With this linearized function for the transcription rate dynamics *F*_*α*_(*t*), the time evolution for *u*_*g*_ and *s*_*g*_ can be decomposed into a sum of effects arising from each module, which gives rise to these new RNA velocity differential equations:

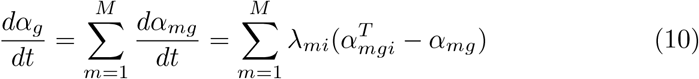

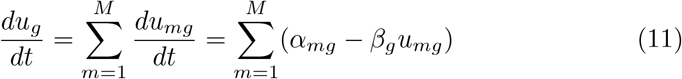

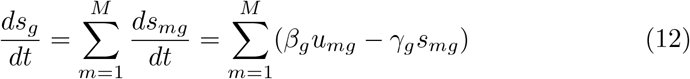

This also defines a velocity for each module, *ν*_*mg*_:

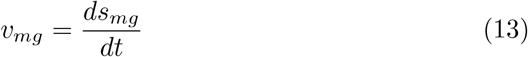

where the total velocity is the sum of the module velocities.

The analytical solution for the transcription rate and spliced and unspliced counts as a function of time is derived in 4.1 and is given by:

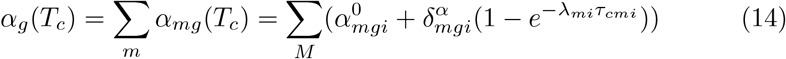

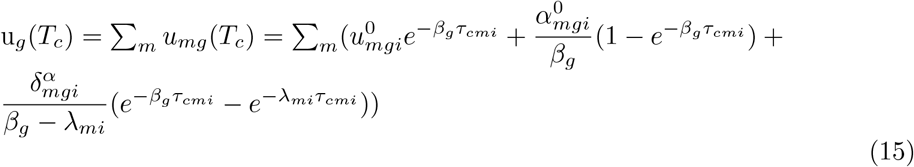

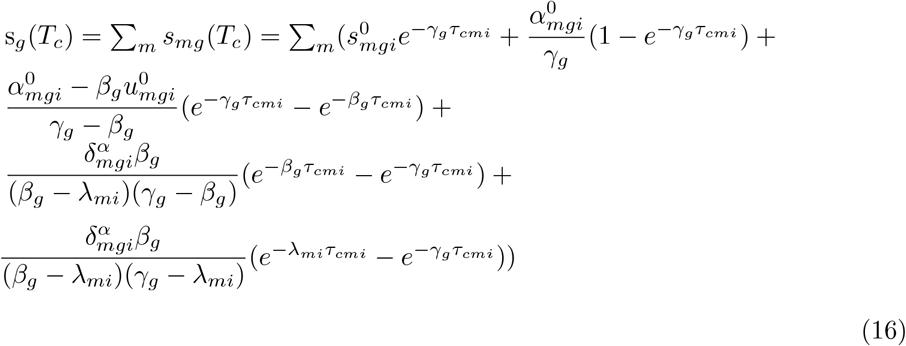

with:

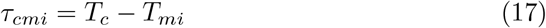

and the following initial conditions:

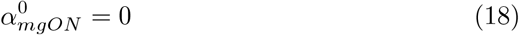

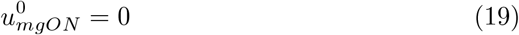

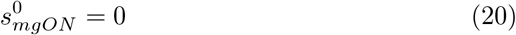

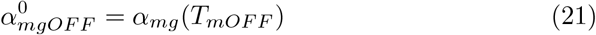

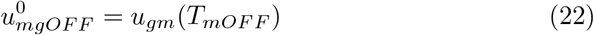

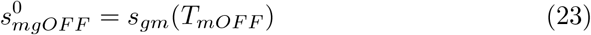

### 1.4 Measurement model

The measurement model of cell2fate is encoded in the likelihood function, which is derived by first transforming “biological” expectation values 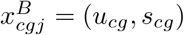 to account for technical variables:

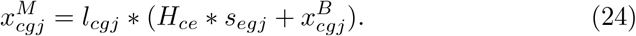

Here, *H*_*ce*_ denotes a one-hot categorical assignment of cells to experimental batches, *l*_*cgj*_ describes differences in detection efficiency (e.g., sequencing depth, read alignment and quantification) of genes across cells and *s*_*gej*_ models ambient RNA (“soup”) for each gene in each batch. These “measurement” expectation values are then used to parameterize a Negative Binomial obsevation model of the observed raw count values *X*_*cgj*_ = (*U*_*cg*_, *S*_*cg*_):

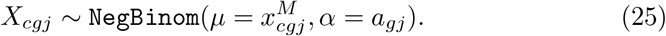

Here, *a*_*jg*_ are Negative Binomial over-dispersion parameters for each gene, separate for spliced and unspliced counts. A Poisson distribution alone can model the sampling noise of transcripts, which is added during measurement [18]. The extra variance added by the Negative Binomial distribution over the Poisson distribution models additional sources of noise [18]. This includes the variance arising from stochastic rates (i.e. “transcriptional bursting”) [18]. While prior work [14, 10] has shown that stochastic rates result in time-varying variances and co-variances for spliced and unspliced counts, we chose not consider this additional complexity for now, because we expect this additional model refinement to have only a small effect on accuracy. Finally, the extra Negative Binomial overdispersion should also account for missing details in our model (i.e. “model uncertainty”), such as the addition of other dynamical processes on top of the main process (e.g. the cell cycle). We chose separate overdispersion parameters for spliced and unspliced counts, since measurement noise is expected to be different for spliced and unspliced counts due to differences in how they are detected and quantified.

**Figure 1:**
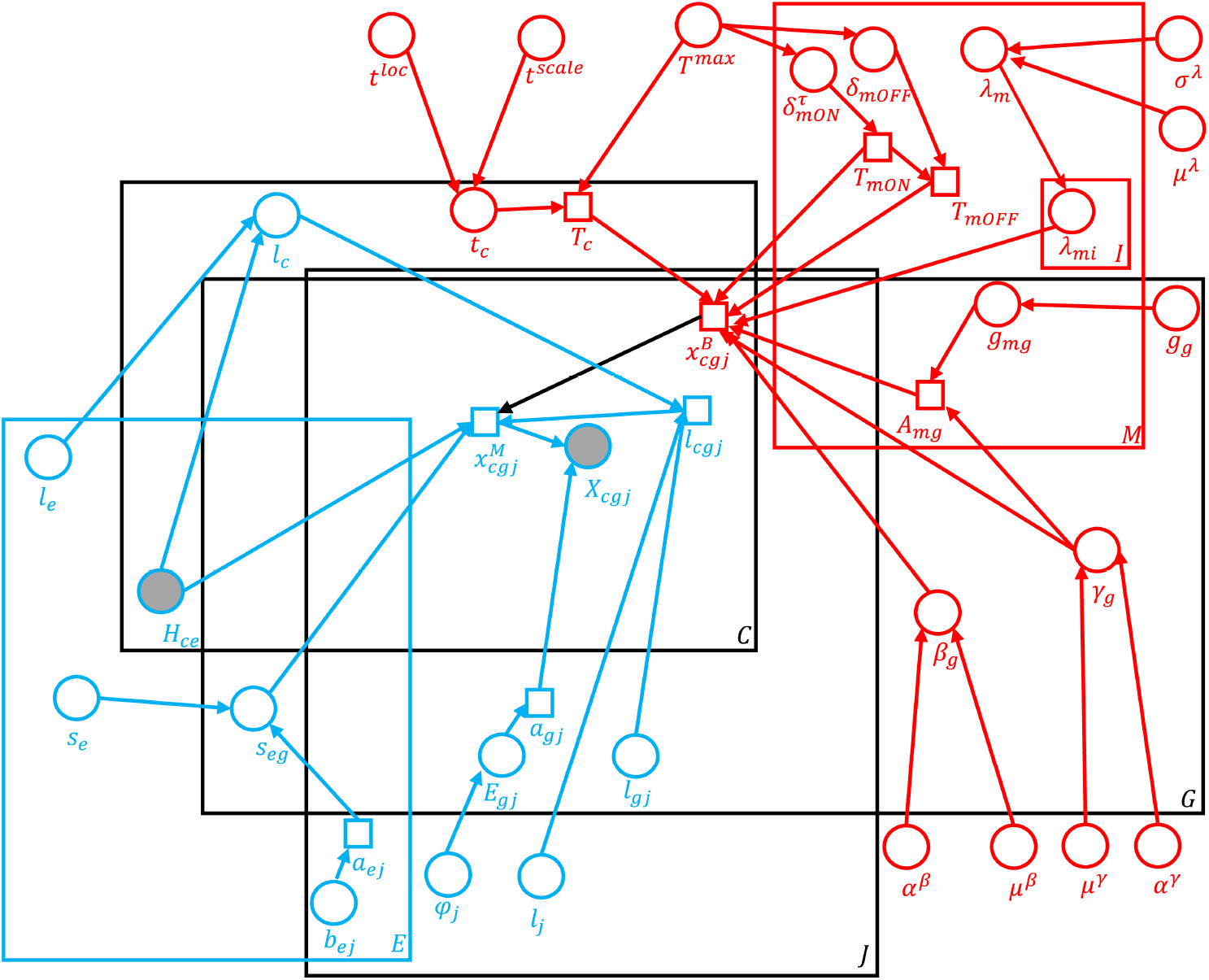
Summary of the generative model in a plate diagram. Circular nodes denote random variables and square notes denote variables that are deterministically computed from other nodes. Red elements are exclusive to the “biological model” and blue elements are exclusive parts of the “measurement model.”

### 1.5 Prior distributions

We derived analytically (Appendix 4.3) that most genes in a module will reach their steady state after 20 hours. This sets a general order of magnitude for the maximal time *T*^*max*^ of a process that we can expect. We choose a prior with large variance at 100 hours, to account for processes with both short and long sequences of module activations:

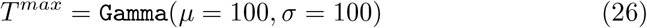

Based on this, the prior for the cell process time is:

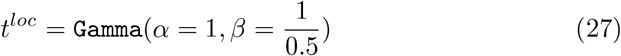

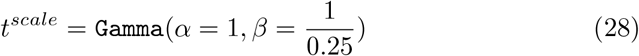

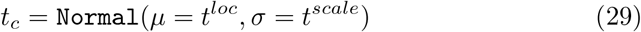

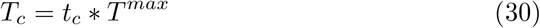

In this parameterization of the Gamma distribution with *α* and *β* instead of *μ* and *σ, α* controls the variance (larger *α* means less variance) and *β* can be set to 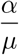 to control the mean. Hence, values for *t*_*c*_ will largely fall between 0 and 1, so that the overall magnitude of *T*_*c*_ is controlled by *T*^*max*^ .

Switch-on times of all modules *m* > 0 are defined by a time gap to the switch-on time of the previous module:

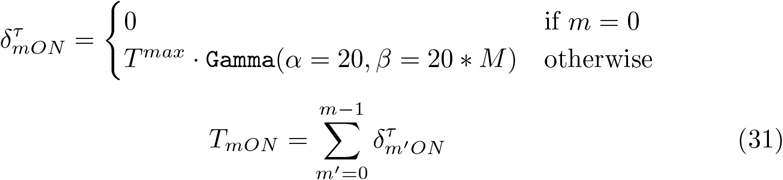

Switch-off times of all modules are assumed to depend on occurrence of a signal with uniform probability of appearing at each point in time, so that an exponential distribution is appropriate:

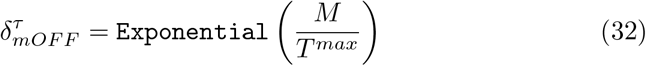

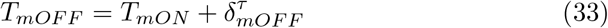

Moving on to the prior distributions for our rate parameters, the mean degradation rate is expected to be 0.2 (molecules/h) and the mean splicing rate is expected to be about 5 times as fast, based on previous work [16]. We chose to set uninformative priors in this general order of magnitude

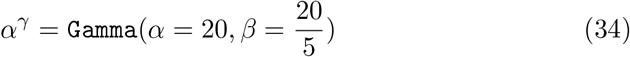

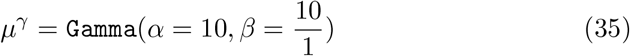

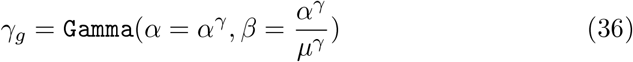

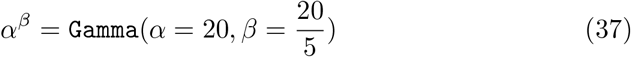

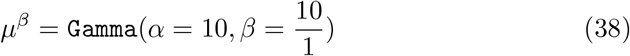

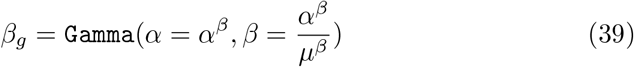

The rates of activation for each module, share a hierarchical prior like this:

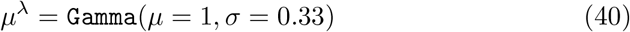

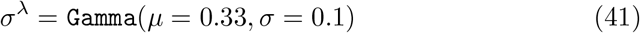

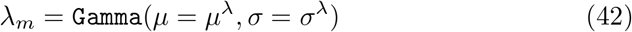

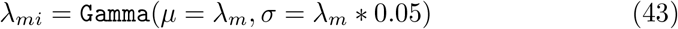

Instead of defining the maximal effect on the transcription rate by each module directly, we define the maximal effect on spliced counts *g*_*mg*_ (at *τ* = ∞) and multiply this by the degradation rate to obtain the effect on the transcription rate:

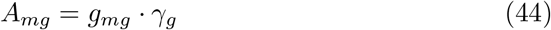

Priors for *g*_*mg*_ are given by:

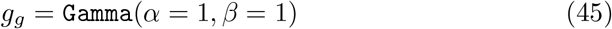

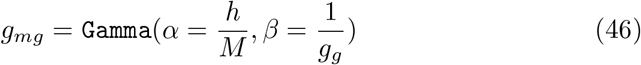

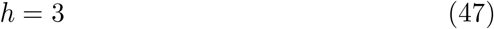

*g*_*g*_ can be interpreted as the total counts produced by a gene in a cell per module in the steady state. *h* can be interpreted as the number of modules this gene is part of, since the mean per module is given by:

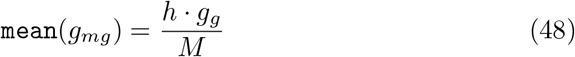

Moving on to technical variables, we use this hierarchical prior for the overdispersion parameter of spliced and unspliced counts:

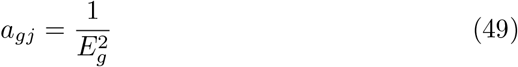

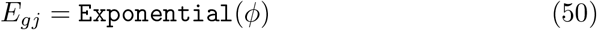

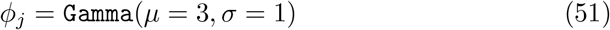

This kind of prior is called a containment prior [20]. To understand the reasoning behind it, it is useful to write out the total variance of the Negative Binomial distribution:

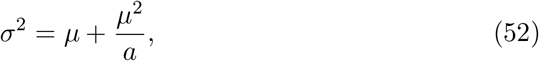

so the chosen prior pushes alpha towards infinity and with that the “extra variance” compared to a Poisson distribution towards 0. So inference should only converge to solutions with considerable overdispersion if this is really necessary.

The mean relative detection probability in each cell depends on the experimental batch *e* each cell comes from:

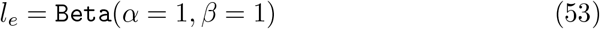

(This results in a broad distribution between 0 and 1 with mean of 0.5.)

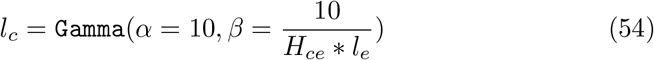

To model differences in detection probabilities for spliced and unspliced counts, we define two more parameters that describe their average difference (using a broad prior around 1):

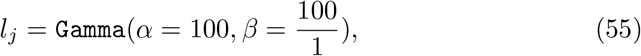

as well as their gene specific difference (using a narrow prior):

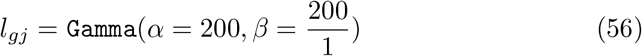

That together give the gene- and cell-specific detection probability for spliced and unspliced counts:

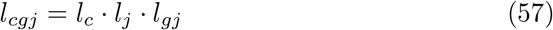

Finally, we use this containment prior for the ambient RNA parameter of both spliced and unspliced counts:

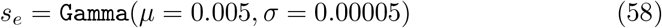

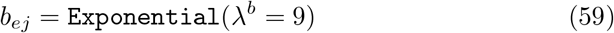

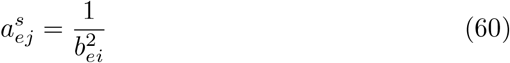

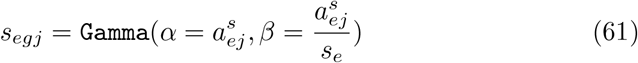

### 1.6 Interpretation of the linearization solution as mixed membership model

The cell2fate model can be parameterized as a mixed membership model, by factoring out the target transcription rate in the on state *A*_*mgON*_ from the time evolution of the total transcription rate (equation 14), like this:

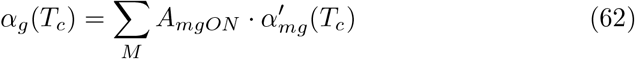

The transcription rate of each module is similar as before:

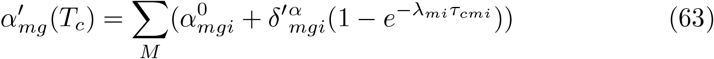

with:

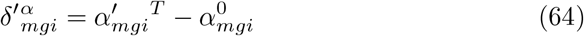

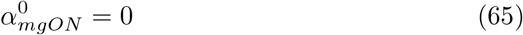

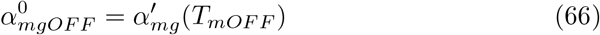

The only difference is that the target transcription rate is now capped at 1:

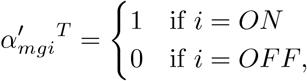

With this parameterization, *A*_*mgON*_ corresponds the “gene loadings” visualized as bars in figure 1B of the main text and 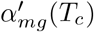 defines the temporal patterns, illustrated at the top of figure 1B, which are similar to “cell loadings” in PCA and non-negative matrix factorization. See appendix 4.4 for further comparisons to non-negative matrix factorization.

### 1.7 Model inference and choice of hyperparameters

The model is implemented in the probabilistic programming language pyro [3]. All parameters are initialized to the mean of their prior distributions. Inference is performed using stochastic variational inference, using the “AutoHierarchical NormalMessenger” autoguide and Adam optimizer [11] for 500 iterations with a learning rate of 0.01 and a batch size of 1,000 cells by default. Posteriors are then estimated numerically by sampling 1000 values from the variational distribution. The parameter controlling the maximal number of modules, *M*, is chosen before training by Louvain clustering of the data and multiplying the total number of clusters by 1.15. For the datasets included in our study this optimization finished in under 20 minutes in all cases (see figure 2).

**Figure 2:**
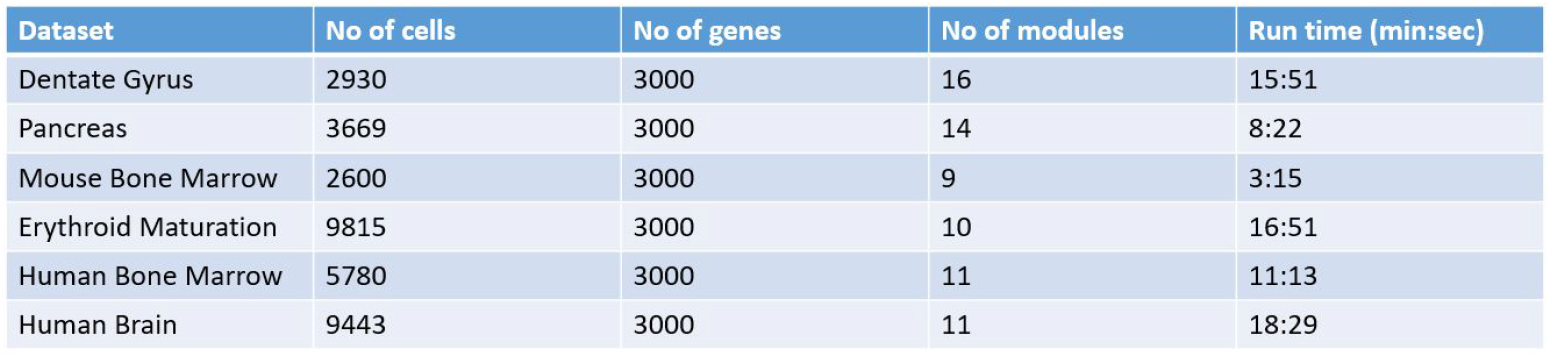
Overview of cell2fate run time on different datasets.

### 1.8 Downstream Analysis

#### 1.8.1 Computation of the RNA velocity graph

We followed the procedure proposed in the scvelo package for computing a cell-cell transition probability graph from RNA velocity estimates [1], with the modification of averaging the velocity graph over 1000 posterior velocity samples, so that noisy gene velocities with high posterior uncertainty have less weight in estimating transition probabilities. Optionally, the original procedure can also be followed exactly, by computing the velocity graph with mean velocity estimates from our method, using the original function in the scvelo python package.

#### 1.8.2 Computation of module activation

We calculate the module activation, which we defined as the total spliced counts produced by a module in a cell, by substituting posterior parameter values into equation 16, which captures the time evolution of spliced counts. Module activation can then be plotted over time, by plotting the posterior time of each cell on the x-axis and the calculated module activation of each module on the y-axis.

#### 1.8.3 Calculation of normalized module activation and state

Module normalized activation denotes the number of counts produced by a module, divided by the steady state counts of this module (figure 3D, grey line), which is calculated by setting time to infinity in equation 16, which captures the time evolution of spliced counts. The module state is defined as OFF if the cell time is smaller than the switch on time of the module or the module normalized activation is below 0.05 (figure 3D of main text, grey) and as ON if its normalized activation is above 0.95 (figure 3D of main text, bright green). Otherwise, a module is in either the induction or repression state depending on whether the inferred cell time is below or above the switch off time (light green or orange in figure 3D of main text).

## 2 Data analysis and experimental methods

### 2.1 Benchmarking of RNA velocity methods

#### 2.1.1 Processing of datasets

We used the 3000 most variable genes with at least 20 detected counts for all results in the manuscript. In addition, for all methods except pyro-velocity and cell2fate we applied the standard preprocessing pipeline suggested in the main scvelo tutorial (on mouse pancreas development). This included total-count normalization, log-transformation and calculating mean expression among the 30 nearest neighbours in 30 component PCA space (“knn-smoothing”).

#### 2.1.2 Application of velocity models

We followed online tutorials based on the mouse pancreas development dataset for all methods and then used the same analysis pipeline to produce the benchmarking results in this manuscript. We summarize parameter settings for all methods in a table in figure 3 and also include the method version, where available. For cell2fate we kept all training and model parameters at their default values, with a single exception: For the Human Bone Marrow dataset we increased the Tmax parameter mean and standard deviation from 50 and 50 to 500 and 100 respectively to cover the larger number of lineages and the longer time-span of human development (as opossed to mouse) covered by this dataset.

#### 2.1.3 Definition of ground truth cell state transitions

For the Mouse Bone Marrow dataset we defined the following ground truth cluster transitions: “dividing” to “progenitors” to “activating”. For the remaining datasets, we used the ground truth transitions from the UniTVelo RNA velocity study that also used these datasets for benchmarking [6]. All ground truth transitions are included in supplementary tables 1 - 5.

#### 2.1.4 Benchmarking metric

We calculated the Cross-boundary directional correctness (CBDir), using the functions provided by the UniTVelo python package [6]. The following is an explanation of the metric copied from the UniTVelo publication [6]: “CBDir measures the correctness of transitions from a source cluster to target cluster using boundary cells given ground truth. Here boundary of source cluster refers to cells in that cluster that are neighbors of target cluster and vice versa. Boundary cells are used because they reflects the biological development in a short period of time and CBDir is calculated via,

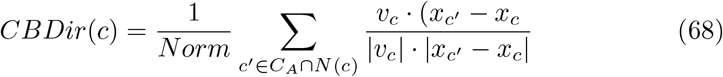

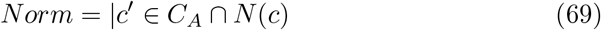

where *C*_*A*_ is sets of cells in target cluster A, *N* (*c*) stands for the neighboring cells of specified cell c. *ν*_*c*_, *x*_*c*_, *x*_*c*_ are the low-dimensional vectors representing computed velocity and positions of cell c and c. Therefore, *x*_*c′*_ −*x*_*c*_ is the displacement is space during the short period of time.”

**Figure 3:**
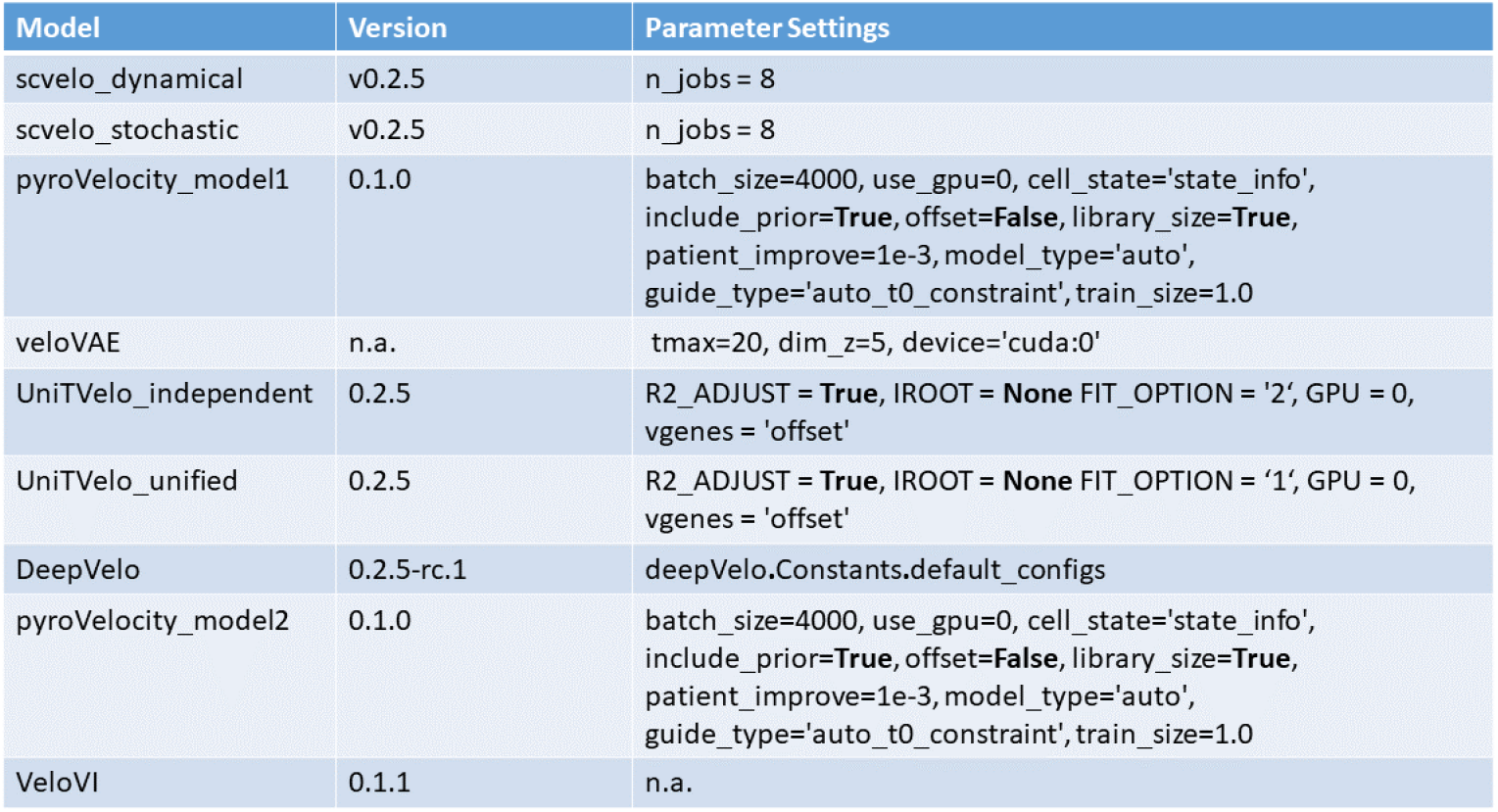
Summary of RNA velocity model versions and parameter settings used in benchmark.

### 2.2 Comparison of decomposition methods on the Dentate Gyrus dataset

#### 2.2.1 Application of cell2fate

Following the benchmarking run on Dentat Gyrus data, we applied the downstream analysis methods, described in sections 1.8.2 and 1.8.3 to calculate and plot module activations and states.

#### 2.2.2 Application of Multi-omics factor analysis (MOFA)

We used the 3000 most variable genes with at least 20 detected counts as input, identical to the cell2fate analysis. We limited the analysis to clusters involved in the neuron differentiation trajectory (Radial glia-like, nIPC, Neuroblast, Granule immature, Granule mature). We added spliced and unspliced counts, normalized, log-transformed and scaled the data matrix using the respective scanpy functions. We ran MOFA using a Gaussian likelihood and 10 factors (the same number of factors found by cell2fate in the relevant clusters). Further run options were: spikeslab_weights = True, ard_factors = True, ard weights = True).

#### 2.2.3 ProBound algorithm

To produce the supplementary figure 18, we applied the probound algorithm [17]. Probound can predict the binding affinity for most transcription factors to a user-supplied DNA sequence. We used the refdata-cellranger-arc-mm10-2020-A-2.0.0 genome as a reference. To get putative promoter sequences for each gene in the reference we extracted the DNA sequence 450 base pairs upstream and 149 base pairs downstream of the transcription start site of each gene. For each transcription factor, we then ranked genes based on the predicted probound binding affinity to its putative transcription start site. We then used the top 10 and bottom 10 binding targets of each transcription factor to produce the results illustrated in supplementary figure 18.

### 2.3 Spatial integration of RNA velocity

#### 2.3.1 Human tissue

Formalin-fixed paraffin-embedded (FFPE) blocks of second trimester human fetal brain were obtained from the Human Developmental Biology Resource, Newcastle, UK (REC 18/NE/0290).

#### 2.3.2 Developing Human Brain snSeq library preparation and sequencing

Single nuclei were isolated from frozen foetal brain tissue per published protocol [12]. Briefly, tissue was Dounce homogenised in homogenisation buffer (250mM sucrose, 25mM potassium chloride, 5mM magnesium chloride, 10mM Tris buffer pH 8.0, 1M 1,4-dithiothreitol, 0.1% Triton X-100, 1X protease inhibitor, 0.4U/L RNAsin Plus RNAse inhibitor, 0.2U/L SUPERaseIn RNAse inhibitor) and filtered with a 40m cell strainer. Debris was removed from the filtrate via density centrifugation with 27All nuclei in a batch were mixed in equal concentrations prior to droplet encapsulation with the 10X Chromium Single Cell 3’ v3.1 kit. Libraries were generated per manufacturer’s protocol (CG000204) and singel-indexed sequenced with cycles 28-8-91 on a NovaSeq 6000 System (Illumina) using a NovaSeq S4 Flowcell.

#### 2.3.3 Developing Human Brain Visium library preparation and sequencing

5*μ*m FFPE tissue sections were stained and imaged following the 10x Genomics Visium CytAssist user guide CG000520. The following times were used for the foetal brain tissue: Haematoxylin 3min, Bluing 1min, Eosin 1min. After probe hybridization and ligation, a Visium CytAssist instrument was used to transfer analytes from the glass slide to a Visium CytAssist Spatial Gene Expression slide with a 42.25 mm2 capture area. The probe extension and library construction were prepared following the standard Visium for FFPE workflow (CG000495) outside of the instrument. Libraries were sequenced with paired-end dual-indexing (28 cycles Read 1, 10 cycles i7, 10 cycles i5, 90 cycles Read 2) in Illumina-HTP NovaSeq 6000 Paired end sequencing in a SP Flow Cell. Loupe browser was used to generate the json file and the Space Ranger pipeline v2022.0705.1 (10x Genomics) and the GRCh38-2020-A reference were used to process FASTQ files.

#### 2.3.4 Developing Human Brain Visium and snSeq count quantification, clustering and annotation

We quantified counts using spaceranger 1.3.0 for the visium data and starsolo with cellranger 3.02 for the single-nucleus data using GRCh38 v1.2.0 as a reference. We applied cellbender to the single-nucleus total count matrix (but not spliced + unspliced count matrices). We followed the scanpy processing and clustering tutorial with default parameters (min genes=200, min cells=3, n genes by counts ¡ 2500, pct counts mt ¡ 5), which involves removing cells and genes with low UMI counts, followed by removal of cells with very high total counts or mitochondrial genes ratio, total-count normalisation, log-transformation, highly-variable gene selection, data scaling, principal-component analysis and finally Louvain clustering. Expression of cell type marker genes taken from the Polioudakis 2019 et al. single-cell atlas figure 1F [**?**] was plotted for each cluster, based on which we annotated the cluster identity. Clusters outside the excitatory neuron lineage (Oligodendrocytes, Interneurons, Microglia) were not considered for further analysis.

#### 2.3.5 Cell2location

We used the steady-state counts of each module as a reference gene expression profile. This steady state expression corresponds to the *g*_*mg*_ parameter of the generative model. We then ran the cell2location method with default parameter settings (including alpha=20).

## 3 Comparative analysis of RNA velocity methods

**Figure 4:**
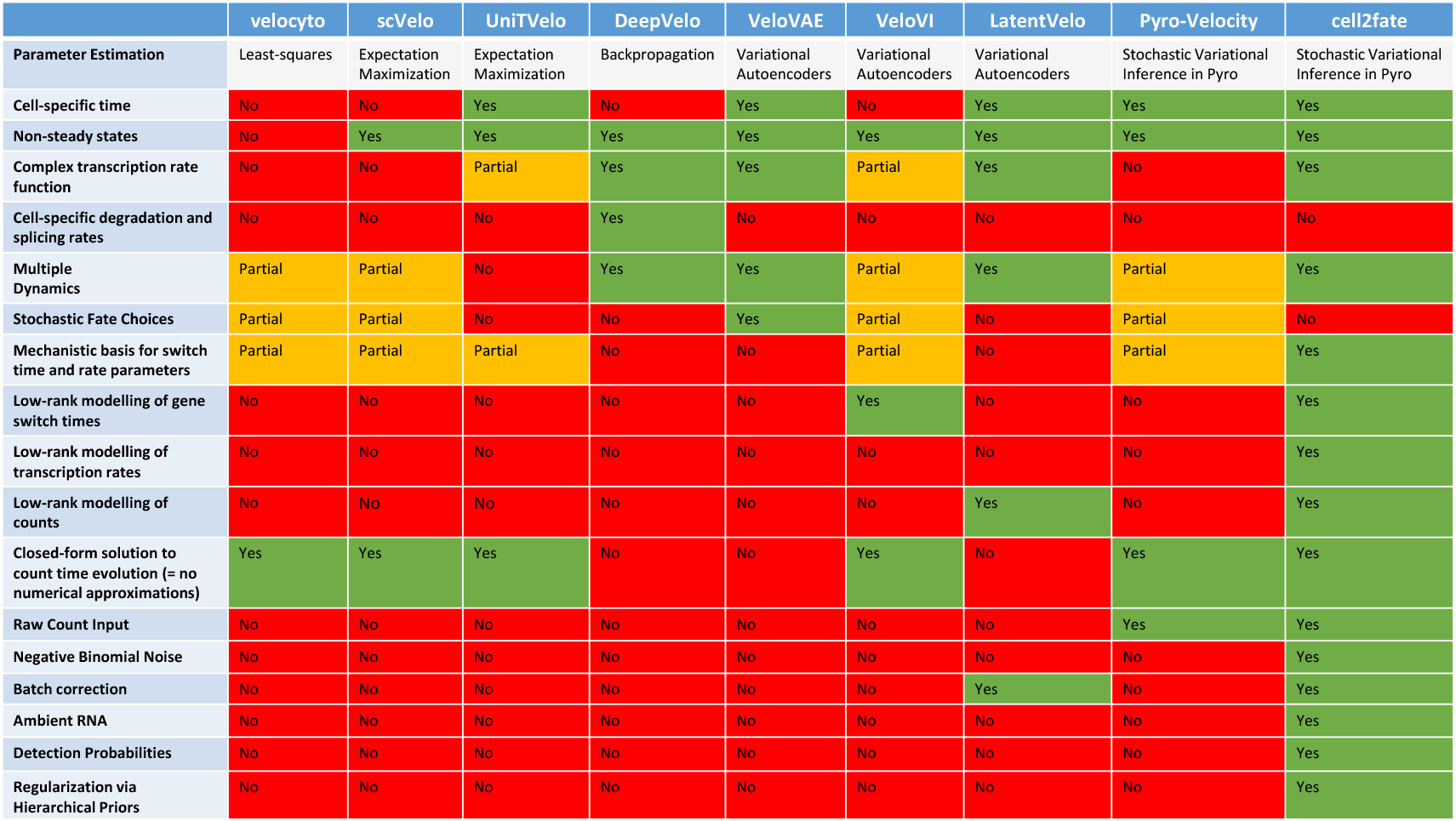
Comparison of RNA velocity methods in table format, according to parameter estimation procedures and model components.

We provide a table in figure 4 that succinctly compares existing RNA velocity methods according to 17 criteria. We further explain and discuss those criteria below.

### 3.1 Cell-specific time

Inuitively, each cell in a dataset can be thought of as occupying a unique process time point in a temporal dimensions, which we called *T*_*c*_ in the cell2fate method. However, velocyto, scVelo and veloVI do not only estimate a separate time for each cell, but also for each gene, resulting in a two-dimensional time parameter *T*_*cg*_. *T*_*cg*_ = 0 is defined as the switch-on time of the gene in this case. In addition, a switch off time *T*_*gOFF*_ is defined separately for each gene. While this approach allows more flexibility in modelling transcriptional dynamics, it increases the number of parameters that need to be estimated by a factor *G* and thus reduces statistical power.

### 3.2 Non-steady states, complex transcription rate function, cell-specific degradation and splicing rates

In general, all methods use the deterministic differential equations describing the transcription, splicing and degradation process, introduced in equations 2 and 3. However, methods differ in how much complexity they allow in the transcription rate function. In addition, DeepVelo stands out as using cell-specific splicing and degradation rates, rather than just gene-specific ones.

The velocyto method solves the RNA velocity differential equations in 2 and 3, by making the assumption that 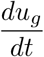 and 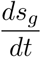 are 0 and then fitting their model to cells with maximal and minimal expression for a given gene. All other methods relax this steady-state assumption and solve the equations in full and then fit their model to all data.

In velocyto, scvelo and pyro-velocity the transcription rate is a step-wise function of time. veloVI and UniTVelo relax this assumption slightly by allowing gradual increases and decreases of the transcription rate, but still with a single maximal value. DeepVelo, VeloVAE, LatentVelo and cell2fate fully relax constraints on transcription rates, by allowing arbitrary transcriptional patterns over time. The way in which the latter four methods achieve this goal is very different. In DeepVelo rate parameters of each cell *α*_*cg*_, *β*_*cg*_, *γ*_*cg*_ are defined as a function of each cell’s gene-expression, *X*_*cg*_ by a graph-convolutional network, *F* :

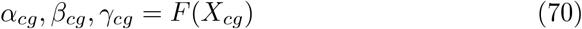

This generates the velocity of each cell, according to the standard equation, as:

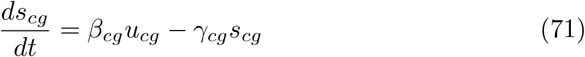

The expected spliced counts expression of each cell is then generated by:

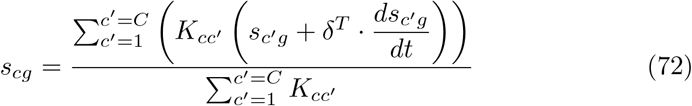

where *δ*^*T*^ is a time-step that is set to 1 in the DeepVelo method and *K*_*cc′*_ is a binary matrix that indicates matching temporal neighbours for each cell that is found as part of the optimization process. The most obvious assumption of this numerical approximation is thus that 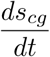 is constant over an interval *δ*^*T*^ = 1. In addition, the approach relays on finding the neighbour matrix *K*_*cc*_ accurately.

VeloVAE has some similarity to the above numerical approach despite being implemented in a different framework, using variational autoencoders. In particular, each cell is assigned a state *c*_*c*_, based on which a relative transcription rate *ρ*_*c*_ between 0 and 1 is generated, by a neural network *F*, so that the total transcription rate is given by:

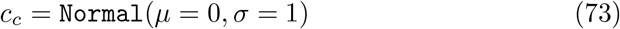

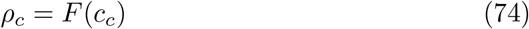

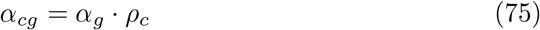

Initial conditions for the differential equations are assumed to be *u*_0_ = *s*_0_ = 0 in a first training run. After the first run, the nearest temporal neighbours of each cell are then used to construct cell specific initial conditions *u*_*c*0_, *s*_*c*0_ for a second training run. Thus this approach assumes transcription rates are constant in the interval between nearest temporal neighbours and it relies on finding those temporal neighbours accurately.

LatenVelo is also based on variational autoencoders, but does not model gene-wise dynamics. Instead it models dynamics in the latent space for unspliced and spliced counts. For this purpose it defines three latent variables:

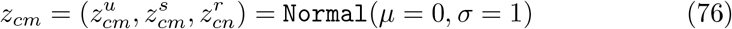

where *m* ∈1, …, *M* and *n* ∈1, …, *N* define the dimensions of latent parameters. *M* = 20 by default and N is adjusted to be the “expected number of lineages” minus 1. 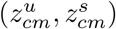 are used to generated expected spliced and unspliced counts, as in a standard variational autoencoder:

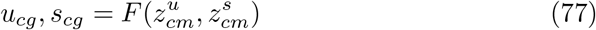

where *F* is the auto-encoder neural network. In addition, all three latent parameters are constrained to obey the following dynamics:

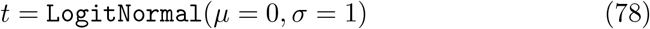

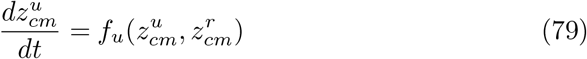

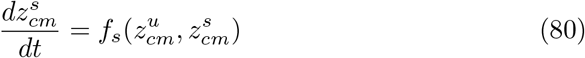

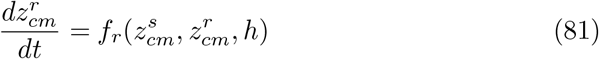

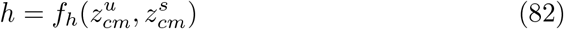

where *f* are neural networks. The rate of change of unspliced counts, thus depends on *z*^*r*^ and *h*, so that this model can in principle capture complex dynamics that depend on the current latent state 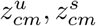 of a cell.

cell2fate describes time-dependent transcription rates, unspliced and spliced counts as the sum over multiple modules effects, as described previously in equations 10 to 12.

### 3.3 Parameter estimation

The original RNAvelocity method velocyto uses least-squares fitting of each parameter. scVelo and UniTVelo deploy maximum likelihood estimation with an Expectation Maximization algorithm. DeepVelo is based on backpropogation. VeloVAE, VeloVI and LatentVelo use backpropogation and mini-batch stochastic gradient descent for variational autoencoders in pytorch, thus enabling posterior parameter uncertainty. Similarly, Pyro-Velocity and cell2fate use stochastic variational inference in pyro.

### 3.4 Multiple dynamics

If an entire group of genes is not expressed at all in one cell type, but expressed in another, UniTVelo can account for such simple differences by assigning a different time to cells in one cell type than another. cell2fate, can more generally account for genes that are transcribed at different non-zero rates in different cell types, by similarly assigning a different time to different cell types, which will result in different modules being turned on in different cell types. Since velocyto, scVelo and veloVI estimate a separate time for each gene, they can also partially account for multiple dynamics. Specifically, a gene that is off in one cell type, but not in the other will simply be assigned a time of 0 for all cells of the latter cell type. However, if a gene is transcribed at different non-zero levels in two cell types, this cannot be accounted for by these models. DeepVelo, LatentVelo and veloVAE can account for multiple dynamics since rates are a function of gene expression or inferred latent state.

### 3.5 Stochastic fate decisions

At branching points the gene expression of a cell is not entirely predictive of its future. Instead, different transcription programs are turned on stochastically. This violates the assumptions of DeepVelo and LatentVelo that make dynamics a deterministic function of current expression. It also violates the assumptions of the cell2fate and UniTVelo methods, that have a transcription rate, as a deterministic function of time. VeloVAE can in principle account for such stochasticity as rates are not a function of gene expression or time, but instead depend on an inferred latent state. Similar to the case of multiple dynamics, the gene-specific time in velocyto, scVelo and veloVI can partially account for stochastic branching points. Specifically, a simple binary decision, where a gene is turned on or not stochastically at a branching point, can be fit with either a non-zero or zero gene-specific time. However, in more complex cases, where a gene stochastically changes its transcription rate to different non-zero values, these models cannot describe the stochastic dynamics accurately. In Pyro-velocity, the switch on point of a gene does not only depend on time, but also on a Bernoulli variable, which can thus capture stochastic decisions, in which a gene is either turned on or not.

### 3.6 Mechanistic basis for switch time and rate parameters

In velocyto, scVelo, veloVI and pyro-velocity genes are assumed to switch on and off at two unique time points. This is only partially realistic, as the transcription rate of each gene, should change continuously in response to abundance of different transcription factors. This is the motivation for cell2fate’s regulatory module formalism. In contrast, in DeepVelo and VeloVI rates change without dependence on a lower-level, mechanistically motivated structure. Finally, in LatentVelo rates are not modelled at all, since latent state velocity and time are inferred directly.

### 3.7 Low-rank modelling of gene switch times, transcription rates or counts

Co-regulation of genes by common transcription factors, results in a low-rank structure to observed dynamics, that can be modelled to increase statistical power. veloVI factorizes the time-matrix *T*_*cg*_ for this purpose, using 10 dimensions by default. LatentVelo represents counts in a latent space of 20 dimensions and models the velocity of these latent states. cell2fate is unique in modelling the fundamental low-rank structure of transcription rates directly. A lower number of switch-times, and a low-rank structure for counts and velocities, results as a downstream result from this modelling choice.

### 3.8 Closed-form solution to count time evolution

velocyto, scvelo, veloVI and pyro-velocity are all based on solving the RNA velocity differential equations analytically, which is possible, because of their simple transcription rate functions. DeepVelo, veloVAE, LatentVelo use numerical approximations to solve their ODEs, which is required given their complex transcription rate functions. In particular, LatentVelo makes use of the “torchdiffeq” package for neural ODEs to compute gradients of inferred latent time points. DeepVelo and veloVAE use approximations as described previously in section 3.2. Cell2fate stands out as having both a complex transcription rate function and an anlalytical solution for the time evolution of spliced and unspliced counts. This is expected to improve performance by circumventing any numerical inaccuracies.

### 3.9 Raw Count Input

pyro-velocity and cell2fate are the only methods that use raw counts as an input. All other methods require preprocessing the data using total-count normalization, log-transformation and nearest-neighbour smoothing. These preprocessing steps are expected to introduce biases for lowly expressed genes (log-transformation) and rare cell-types (nearest-neighbour smoothing).

### 3.10 Negative Binomial Noise, Batch Correction, Ambient RNA, Detection Probabilities

Both LatentVelo and cell2fate include a batch variable in their model. Cell2fate stands out as the only method to model ambient RNA, detection probabilities and Negative Binomial overdispersion (all seperate for spliced and unspliced counts).

### 3.11 Regularization via Hierarchical Priors

Cell2fate makes use of the flexibility of the pyro-probabilistic programming language by using hierarchical priors for all cell and gene specific parameters, which is expected to provide parameter regularization in a principled manner.

## 4 Appendix

### 4.1 Derivation of transcription rate, spliced and unspliced counts as a function of time

We drop both subscripts and the summation over all modules for simplicity. Starting from our differential equation for the transcription rate:

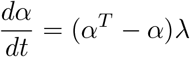

This can be solved directly by integrating:

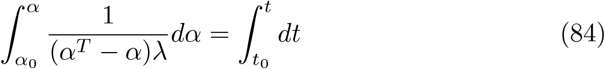

To yield:

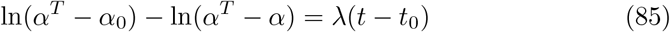

Defining:

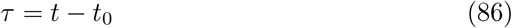

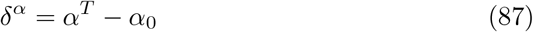

And rearranging terms we obtain our solution for *α* :

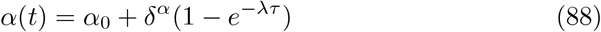

Substituting into the equation for unspliced counts:

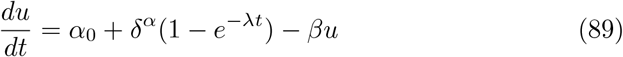

We multiply both sides by *e*^*βt*^ and solve this via integration:

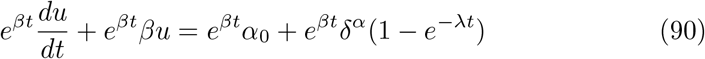

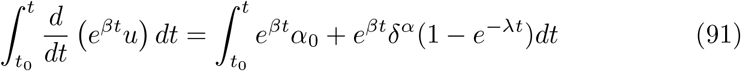

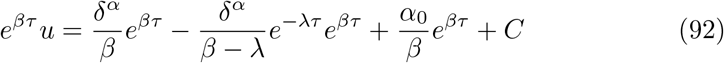

C is given by substituting the initial condition (*u*_0_, *t*_0_):

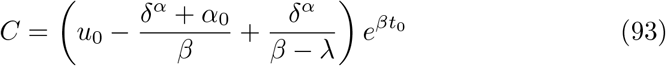

Substituting C and rearranging terms gives us our equation for unspliced counts:

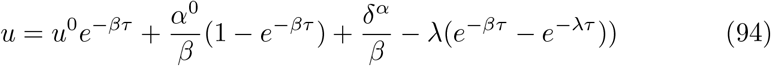

The solution for spliced counts is obtained in a similar manner. We first substitute our expression for unspliced counts, then multiply by *e*^*γt*^ and then integrate:

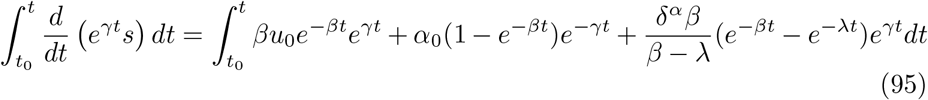

to obtain:

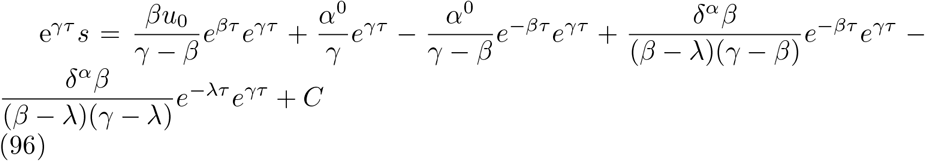

The initial condition is found at the point (*t*_0_, *s*_0_) and after rearranging terms we obtain the solution for spliced counts.

### 4.2 Effect of mRNA detection probabilities on observed dynamics

We define observed spliced and unspliced counts as *s*^*obs*^ = *l*_*s*_ *· s* and *u*^*obs*^ = *l*_*u*_ *· u* respectively, i.e. *l*_*s*_ and *l*_*u*_ are detection probabilities for spliced and unspliced counts respectively.

For simplicity we make the approximation that *λ >>* 1. This means changes in the transcription rate do not happen gradually over time, but instead occur instantaneously. This means we can use the simpler equations from the scvelo model [1] and now multiply both sides by *l*_*u*_ and *l*_*s*_ respectively:

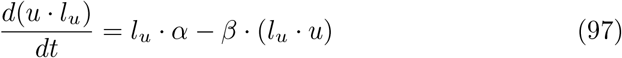

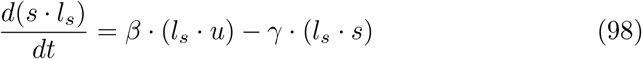

Substituting expressions for observed counts we obtain:

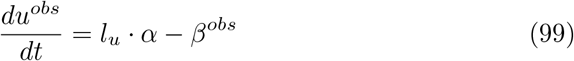

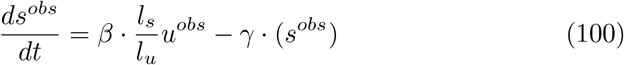

This shows the importance of modelling detection probabilities. Otherwise, observed dynamics for spliced counts will appear to have a different splicing rate than unspliced counts by a factor of 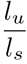 and the transcription rate will appear smaller by a factor *l*_*u*_. This also implies that modelling relative detection probabilities between spliced and unspliced counts is enough, if we do not care about the absolute magnitude of the transcription rate for downstream analysis. Total-count normalization cannot remove this effect, since total counts can vary for biological reasons and not just due to detection probabilities.

### 4.3 Approximate time until steady-state spliced counts abundance

We aim to find the time at which spliced counts reach *r* = 0.99 of their maximal abundance, so 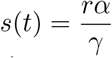 .

We assume *λ >>* 1 so that we can use this simple form for spliced counts, first suggested in the scvelo model [1]:

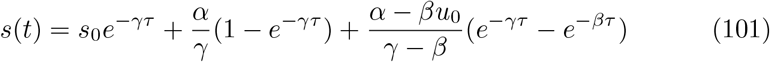

For most genes, *β > γ* [16], so that at *τ >>* 1, exponential terms with *γ* will dominate the spliced counts equation:

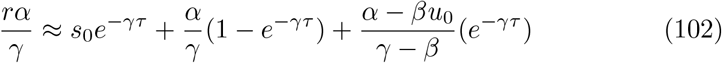

Rearranging for *τ* and assuming *u*_0_ = *s*_0_ = 0:

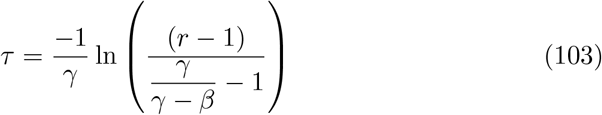

And substituting typical values of *β* = 1 mol*/*h and *γ* = 0.2 mol/h :

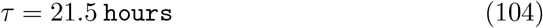

This can be seen as a general order of magnitude at which most genes will reach most of their maximal abundance. However, some genes will be much faster and others much slower, depending on their splicing and degradation rates. If we do not assume *u*_0_ = *s*_0_ = 0 and *λ >>* 1 this time will also depend on the *α* and *λ* values for each gene.

### 4.4 Connection to non-negative matrix factorization

There is an interesting connection of our model to non-negative matrix factorization (NMF). In principle, the *g*_*mg*_ values could be found with a simple NMF model, defined as:

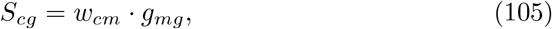

where *w*_*cm*_ would take on different values between 0 and 1 depending on how close a cell is to the steady state for the module. For example, this prior could be appropriate:

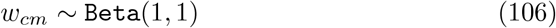

However, as we show in 4.3, how quickly a gene approaches its steady-state depends on the various rate parameters of a gene. So summarizing the similarity to the steady state with a single gene-shared parameter *w*_*cm*_ effectively assumes that all genes have the same rate parameters, which is unrealistic. Hence, NMF will generally find a different set of *g*_*mg*_ values. This also shows the problem with using dimensionality reduction methods as a preprocessing step as typically done in “knn-smoothing”.

### 4.5 Extensions

Since the cell2fate model is under continuous development, we describe here three current development directions to make it easier for the community to contribute on the cell2fate github repository.

#### 4.5.1 Extension to stochastic rates

Stochastic rates have two important effects on our system. Firstly, the mean transcription rate of each gene, cannot be assumed to follow a single deterministic path as a function of time and instead will split into two paths at each lineage bifurcation point. Secondly, the overdispersion parameter of the Negative Binomial distribution should vary with time [14, 10].

An approximate solution to model the first effect, would be to model module activation times as a Markov Process. For this purpose, we define an index *t* with maximal value *T* = *M* to denote transitions in the Markov Chain. Furthermore, we define a new index *n* with maximal value *N* to denote the states of the Markov chain. The current Markov state of a cell is defined as its last activated module. We define *N* = *M* + 2 to include an initial state *N−*1 with no past activated modules and a final state *N* with no future activated modules. This means transition probabilities between states are given by:

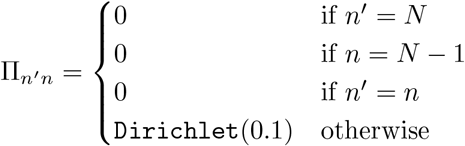

For each cell the Markov sequence of states is then given by:

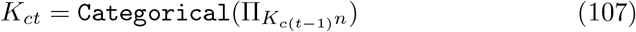

Analogous to equation 111, activation times would then be given by:

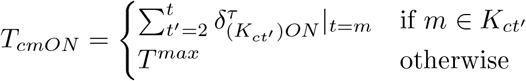

#### 4.5.2 Extension to variable splicing and degradation rates

An approximate solutions to remove limitation 2.) and to introduce time-varying splicing and degradation rates would be to have different rates for each module and state. For example, we could change *β*_*m*_ to *β*_*mgi*_, so that for unspliced counts we have:

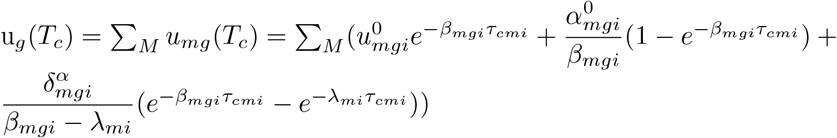

Since each module-specific splicing rate would only apply to counts produced by that module, the splicing rate for each gene at each point in time would effectively be an average, weighted by the counts of each module at each point in time.

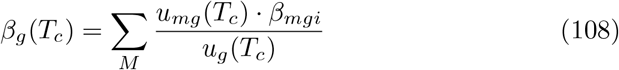

Time-varying degradation rates would follow a similar reasoning.

#### 4.5.3 Extension to link regulatory genes and transcription rate effects

Currently, the effect on transcription rates by each module *A*_*mg*_ is given by a prior distribution that does not share information across time steps (i.e. equations 44 to 46). Instead, we could aim to infer a non-linear function that makes the transcription rate effect of module *m* the result of the *l* previous module transcription rate effects. For example, *l* = 1 would consider just the last module:

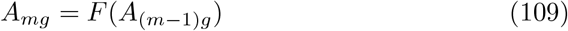

In addition, we could introduce the constrain that only one module is active at each point in time, by using switch times of this form:

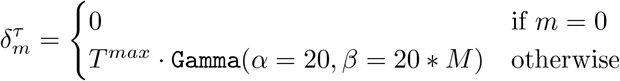

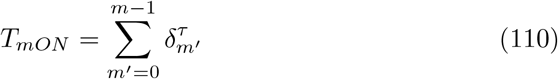

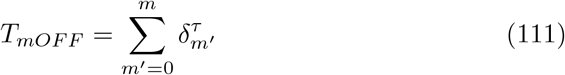

This approach can be interpreted as modelling the effects of the specific regulatory protein abundance, that is present at each discrete time step *δ*^*τ*^ on the next time step.

### 4.6 List of all components in the cell2fate generative model

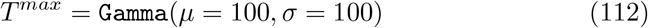

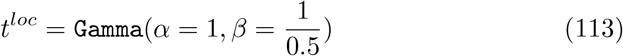

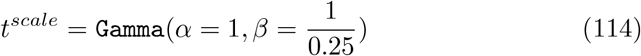

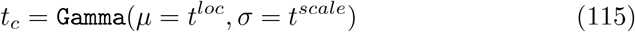

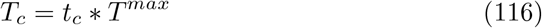

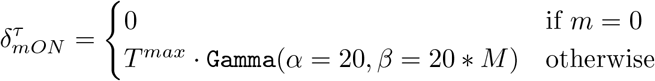

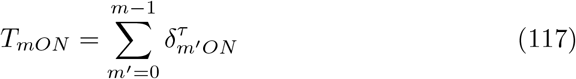

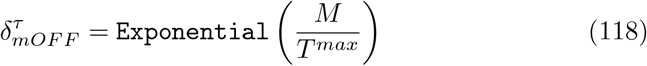

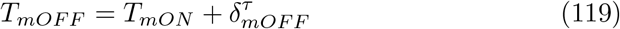

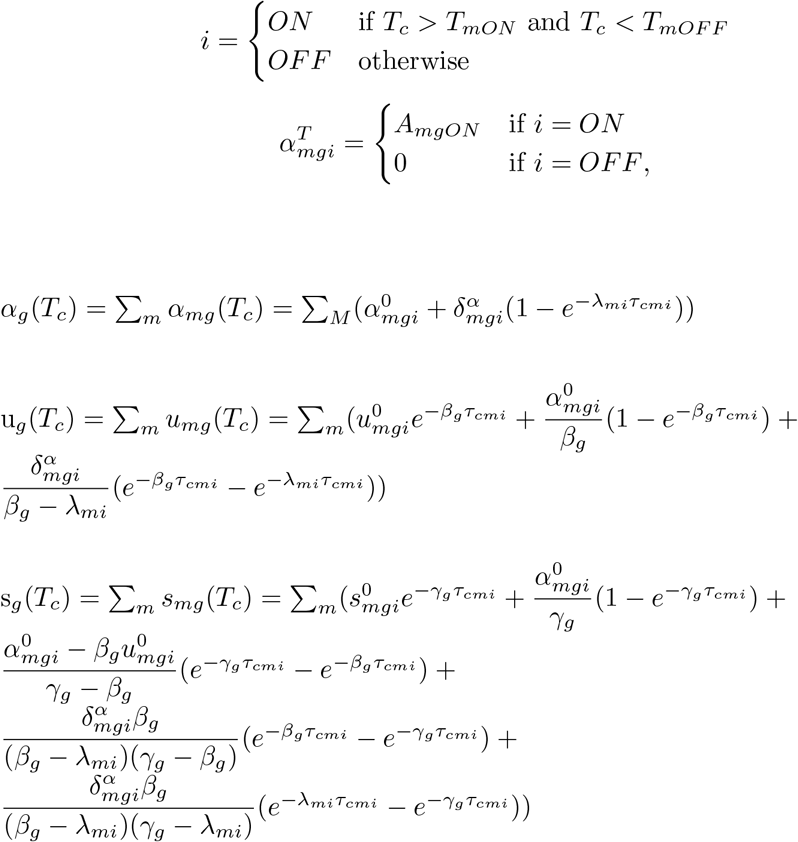

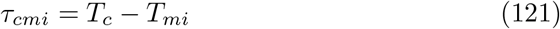

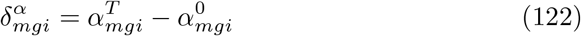

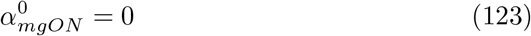

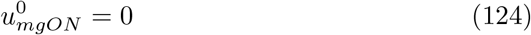

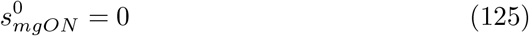

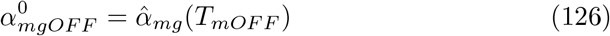

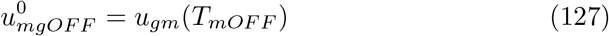

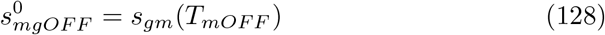

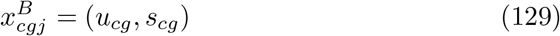

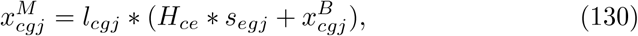

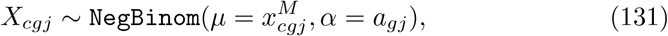

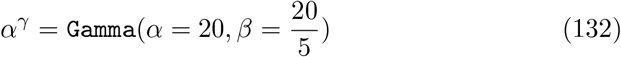

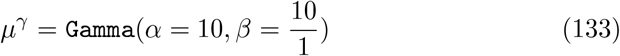

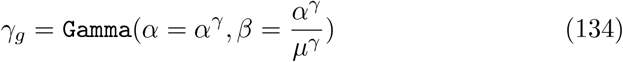

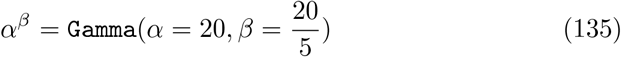

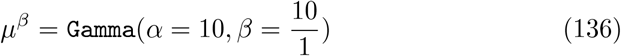

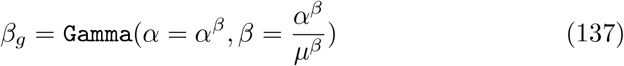

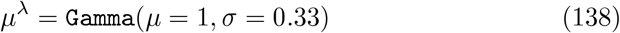

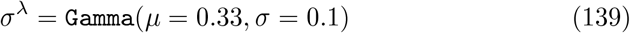

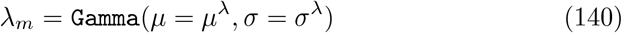

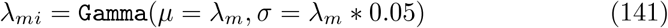

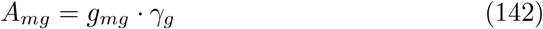

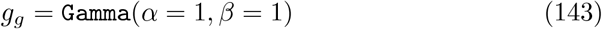

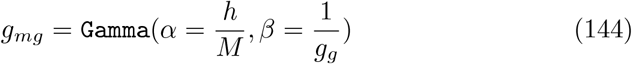

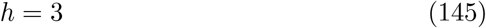

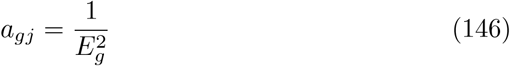

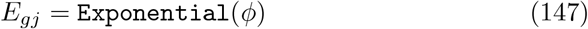

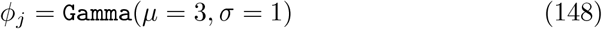

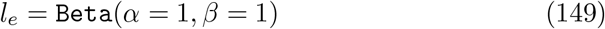

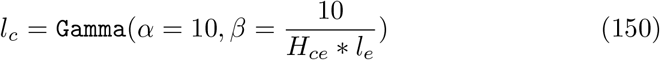

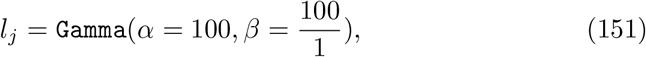

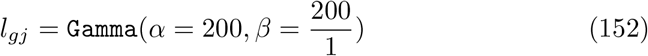

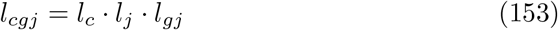

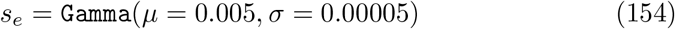

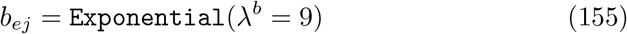

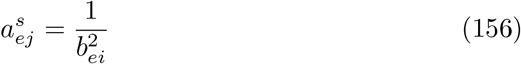

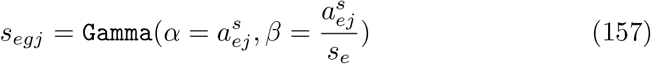

