## SupplementaryFigures for "Model-based inference of RNA velocity modules improves cell fate prediction"

### Supplementary Figures

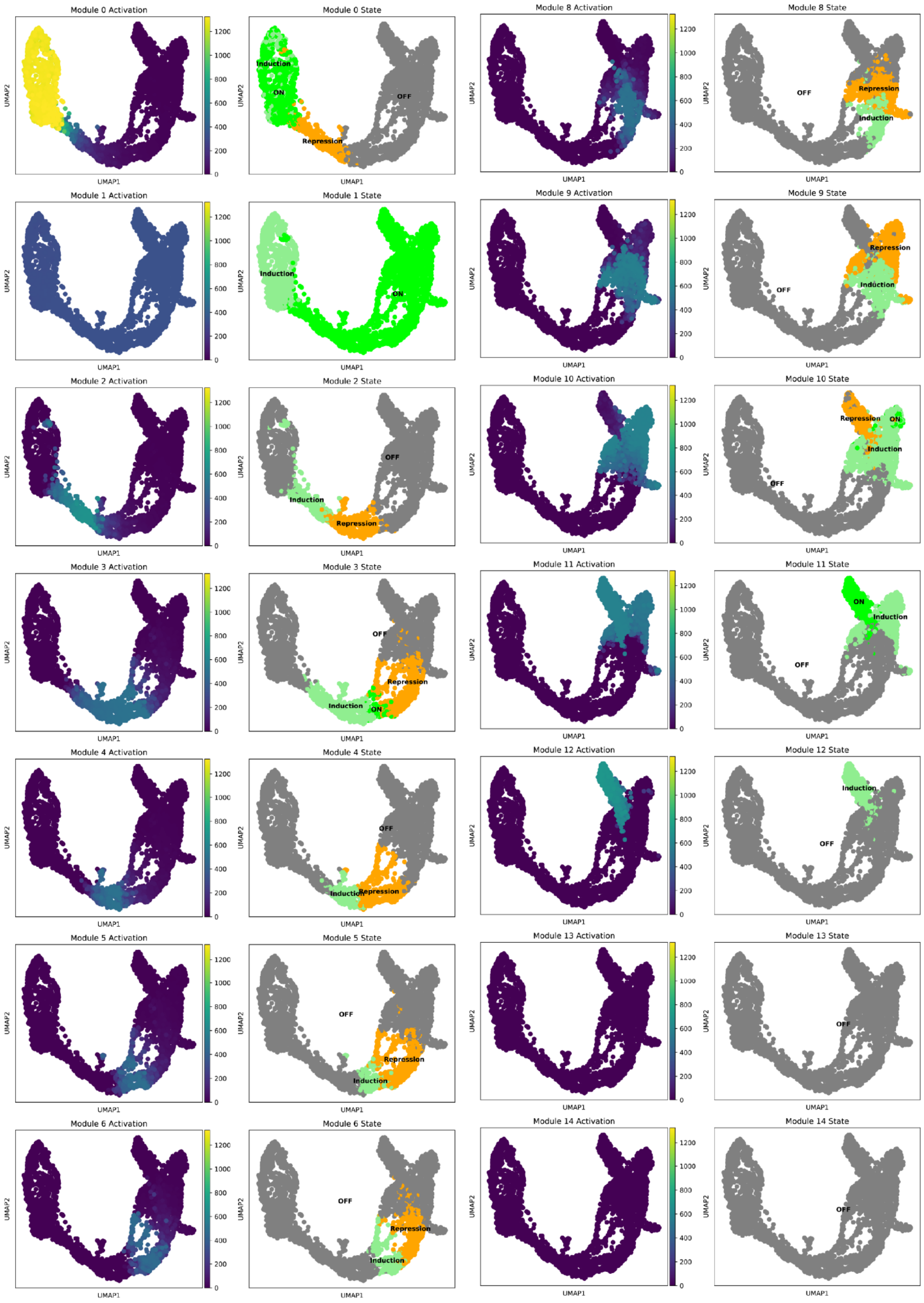


**Supp. Figure 1: module activation and state UMAP plots for mouse pancreas data**
*UMAPs coloured by module activation in columns 1 and 3 and state in columns 2 and 4, highlight a sequence of transcriptional programs switched on during differentiation.*

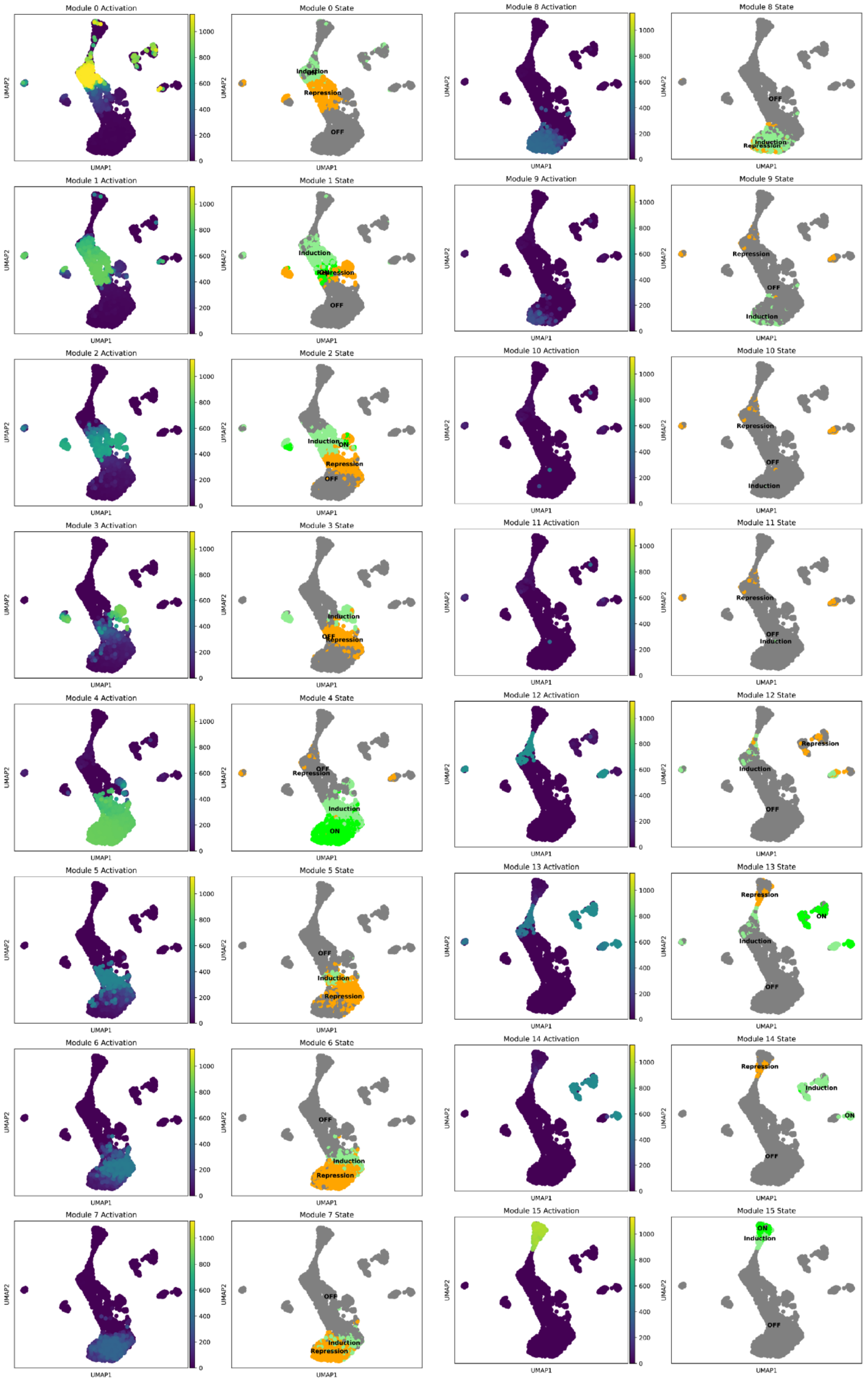


**Supp. Figure 2: module activation and state UMAP plots for mouse dentate gyrus data**

*UMAPs coloured by module activation in columns 1 and 3 and state in columns 2 and 4, highlight a sequence of transcriptional programs switched on during differentiation.*

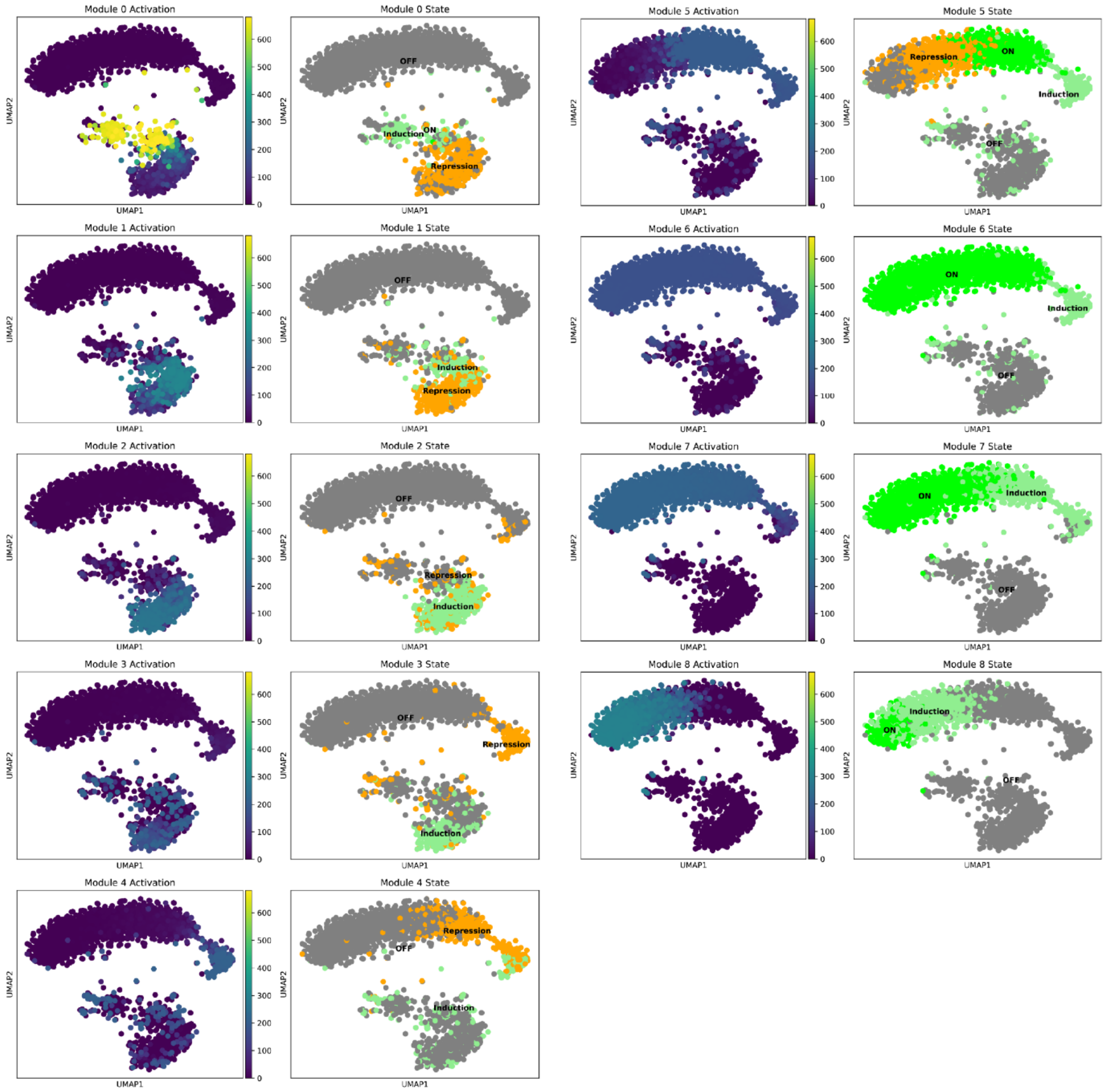


**Supp. Figure 3: module activation and state UMAP plots for mouse bone marrow data**

*UMAPs coloured by module activation in columns 1 and 3 and state in columns 2 and 4, highlight a sequence of transcriptional programs switched on during differentiation.*

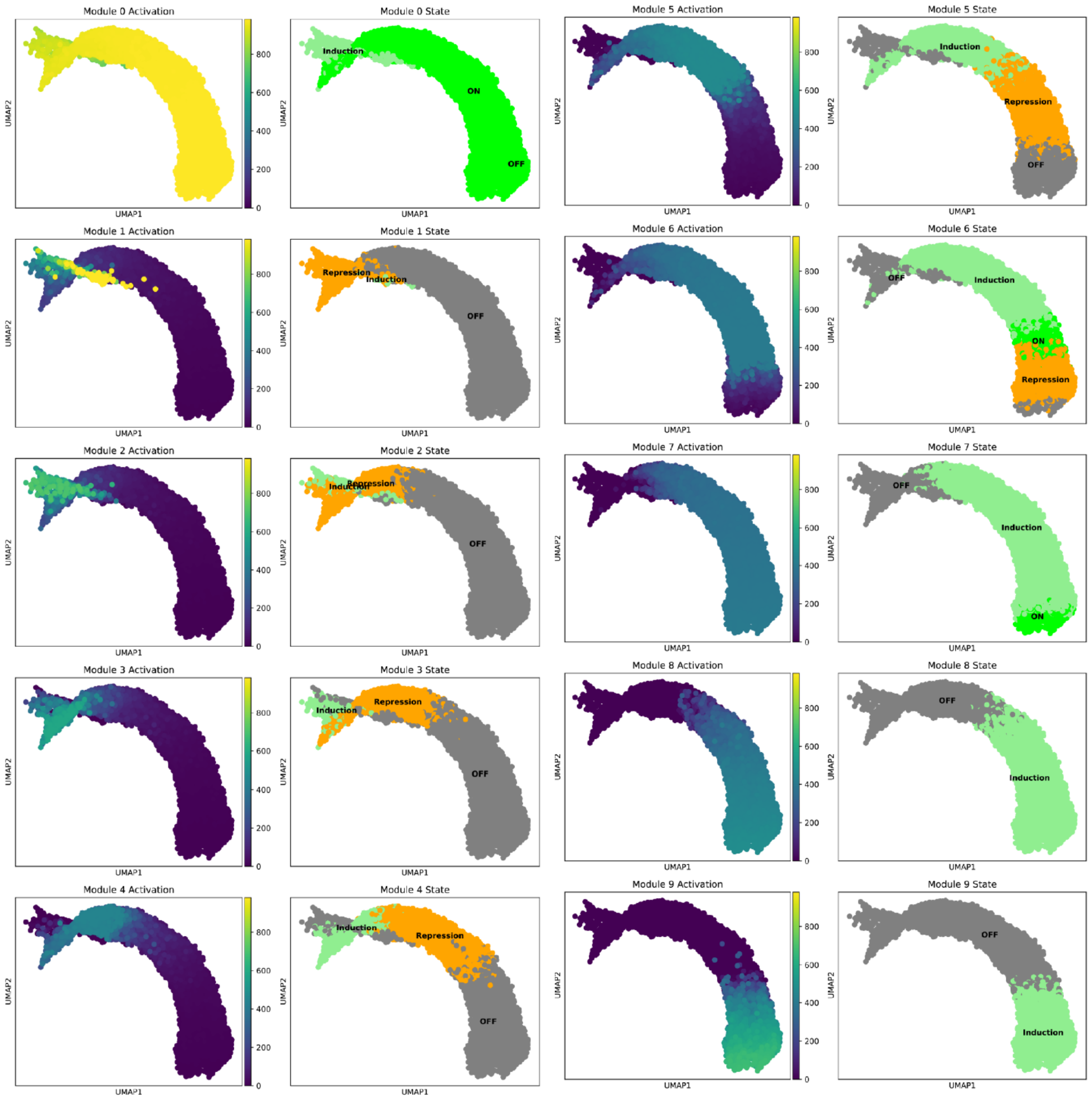


**Supp. Figure 4: module activation and state UMAP plots for mouse erythroid maturation**

*UMAPs coloured by module activation in columns 1 and 3 and state in columns 2 and 4, highlight a sequence of transcriptional programs switched on during differentiation.*

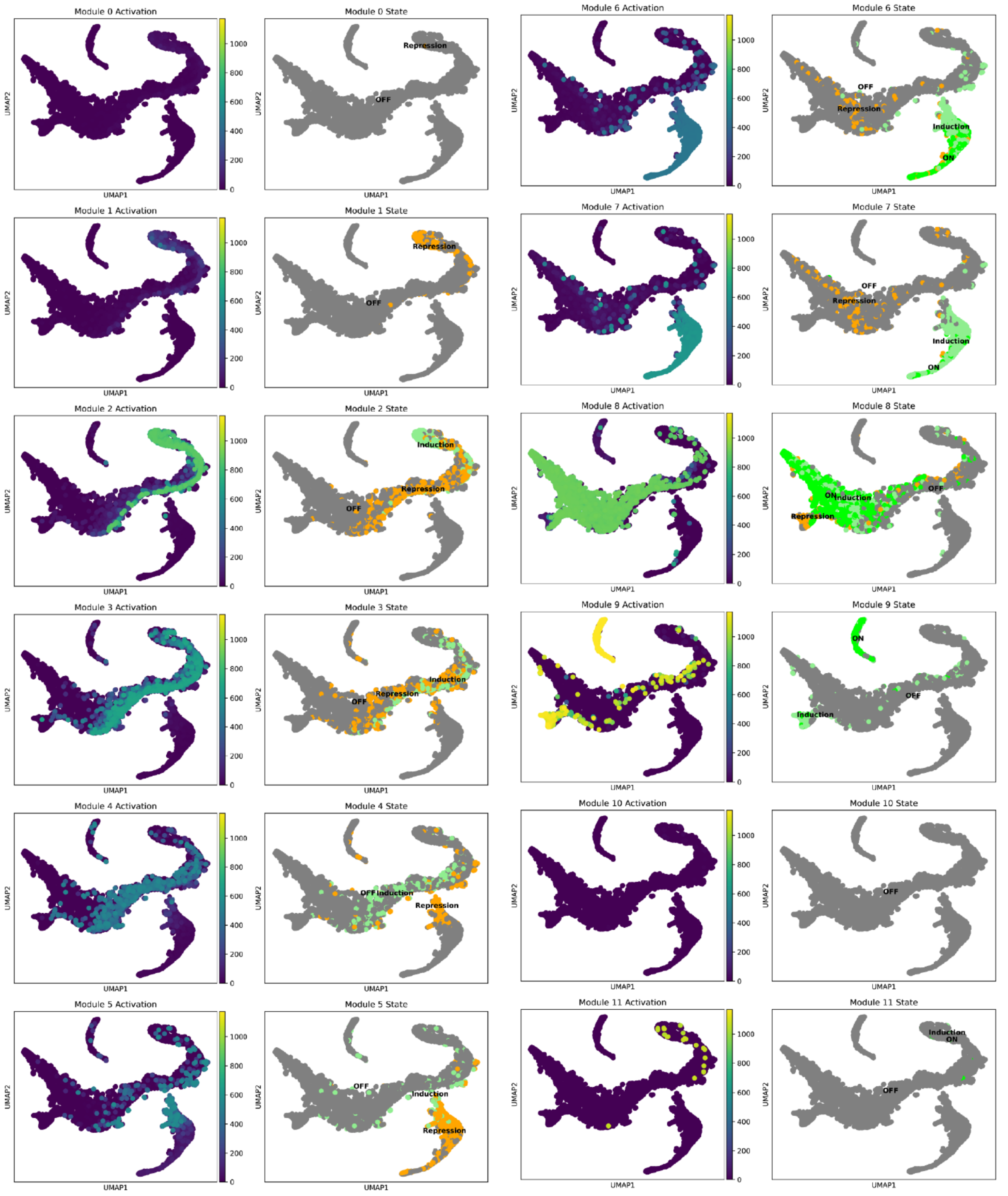


**Supp. Figure 5 : module activation and state UMAP plots for human bone marrow data**

*UMAPs coloured by module activation in columns 1 and 3 and state in columns 2 and 4, highlight a sequence of transcriptional programs switched on during differentiation.*

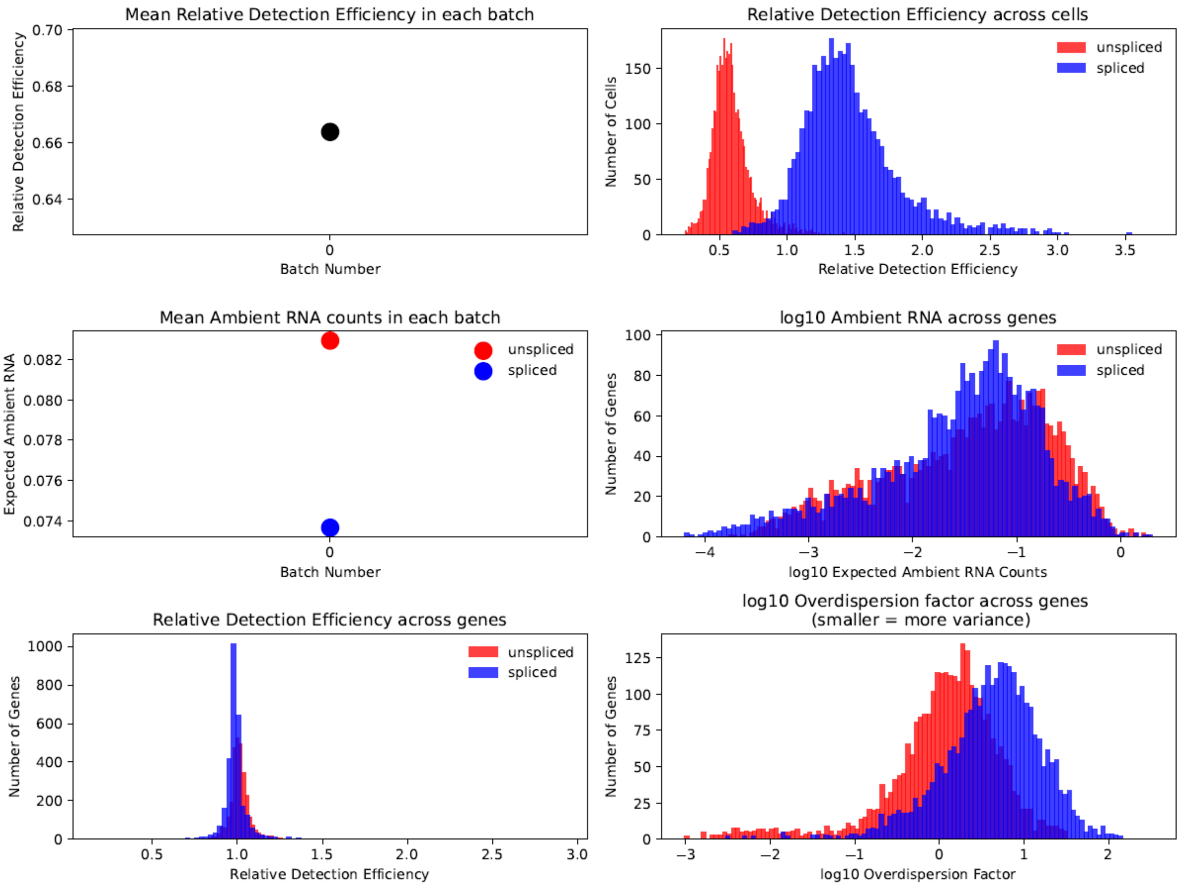


**Supp. Figure 6: overview of noise and technical variables for mouse pancreas data**

*Unspliced counts have lower detection efficiency (top right), higher ambient RNA (middle right) and more noise than spliced counts, corresponding to a lower overdispersion parameter (lower right).*

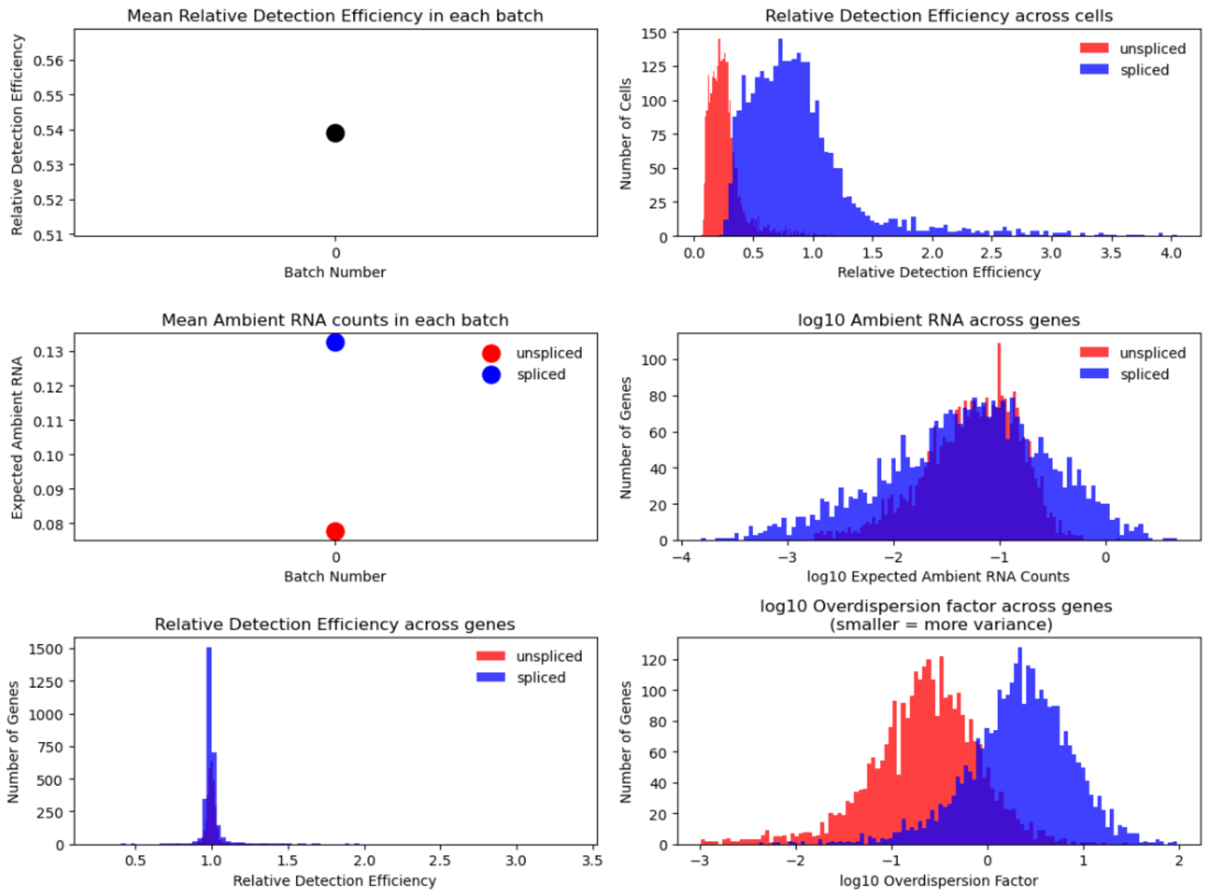


**Supp. Figure 7: overview of noise and technical variables for mouse dentate gyrus data**
*Unspliced counts have lower detection efficiency (top right), higher ambient RNA (middle right) and more noise than spliced counts, corresponding to a lower overdispersion parameter (lower right).*

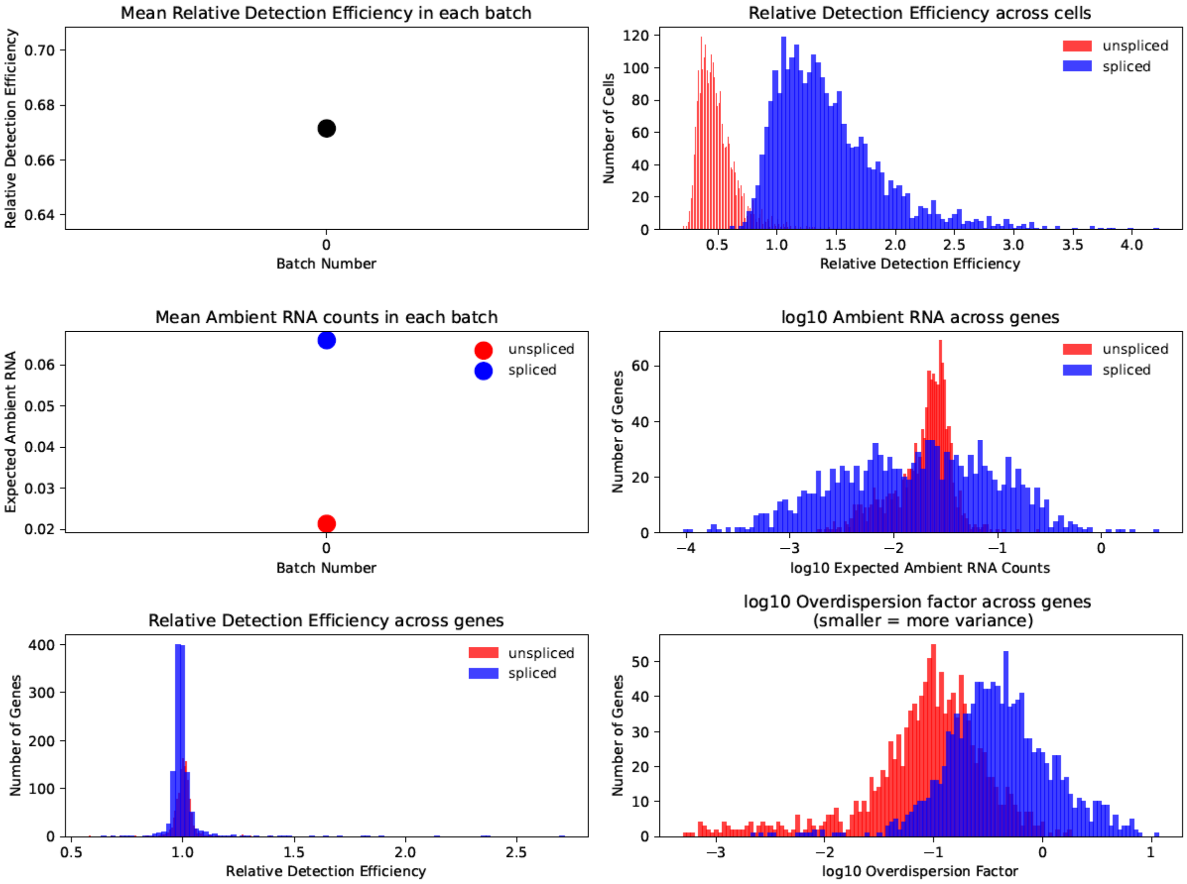


**Supp. Figure 8: overview of noise and technical variables for mouse bone marrow data**

*Unspliced counts have lower detection efficiency (top right), higher ambient RNA (middle right) and more noise than spliced counts, corresponding to a lower overdispersion parameter (lower right).*

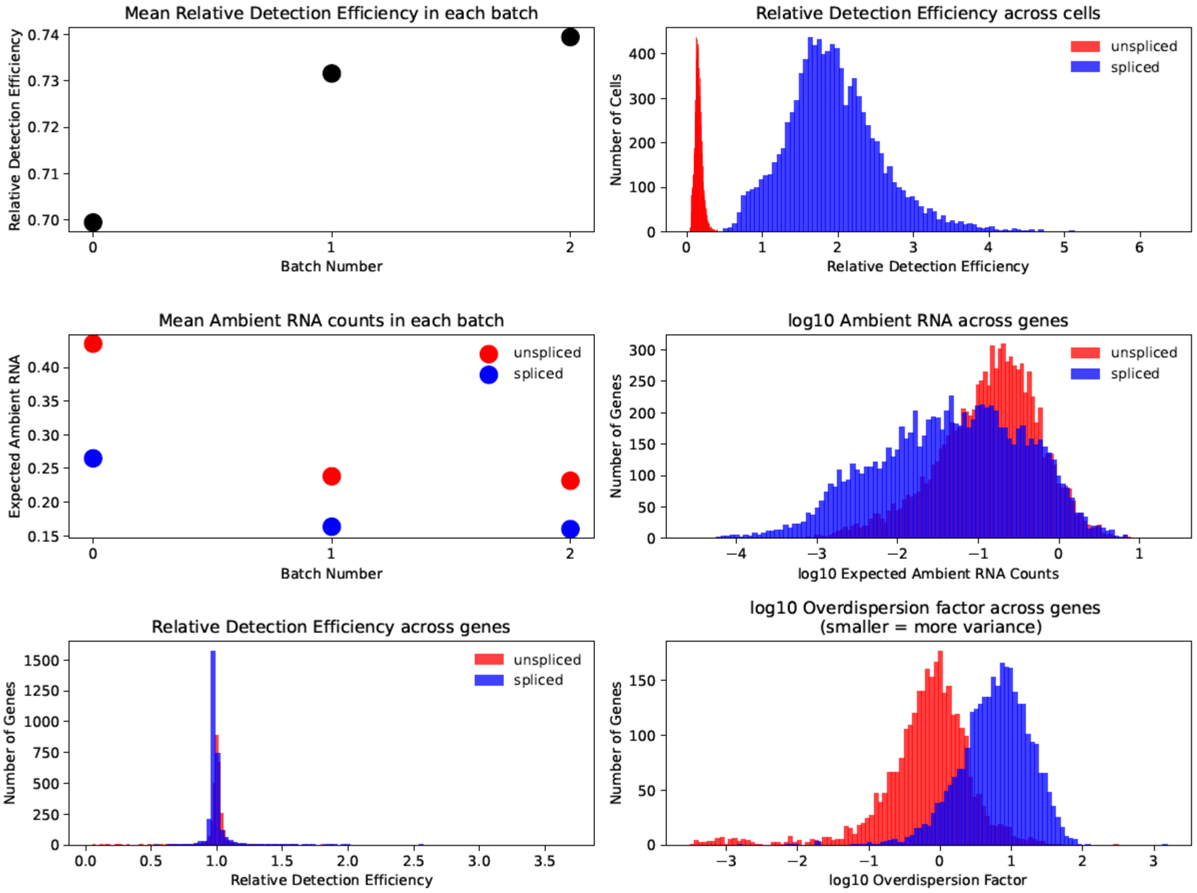


**Supp. Figure 9: overview of noise and technical variables for mouse erythroid maturation data**

*Unspliced counts have lower detection efficiency (top right), higher ambient RNA (middle right) and more noise than spliced counts, corresponding to a lower overdispersion parameter (lower right).*

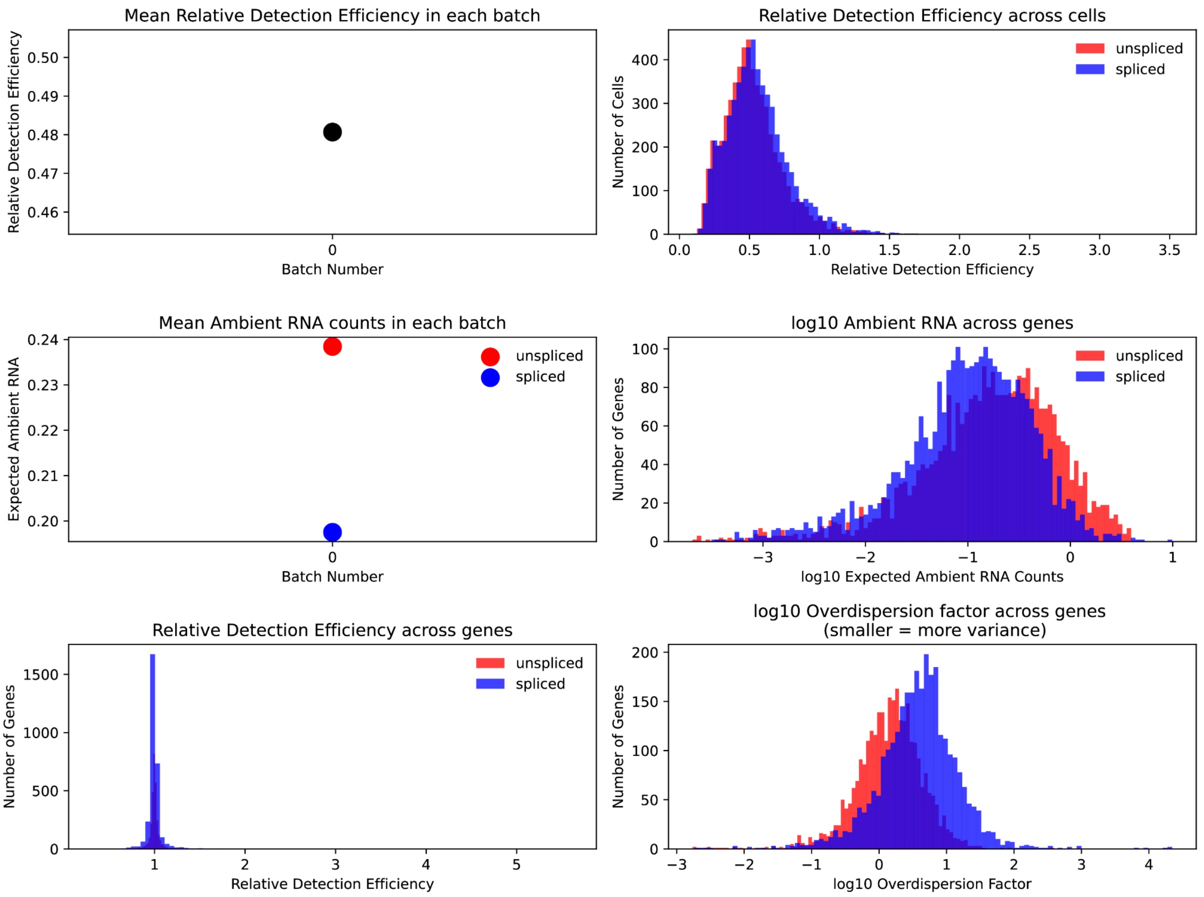


**Supp. Figure 10: overview of noise and technical variables for human bone marrow data**

*Unspliced counts have lower detection efficiency (top right), higher ambient RNA (middle right) and more noise than spliced counts, corresponding to a lower overdispersion parameter (lower right).*

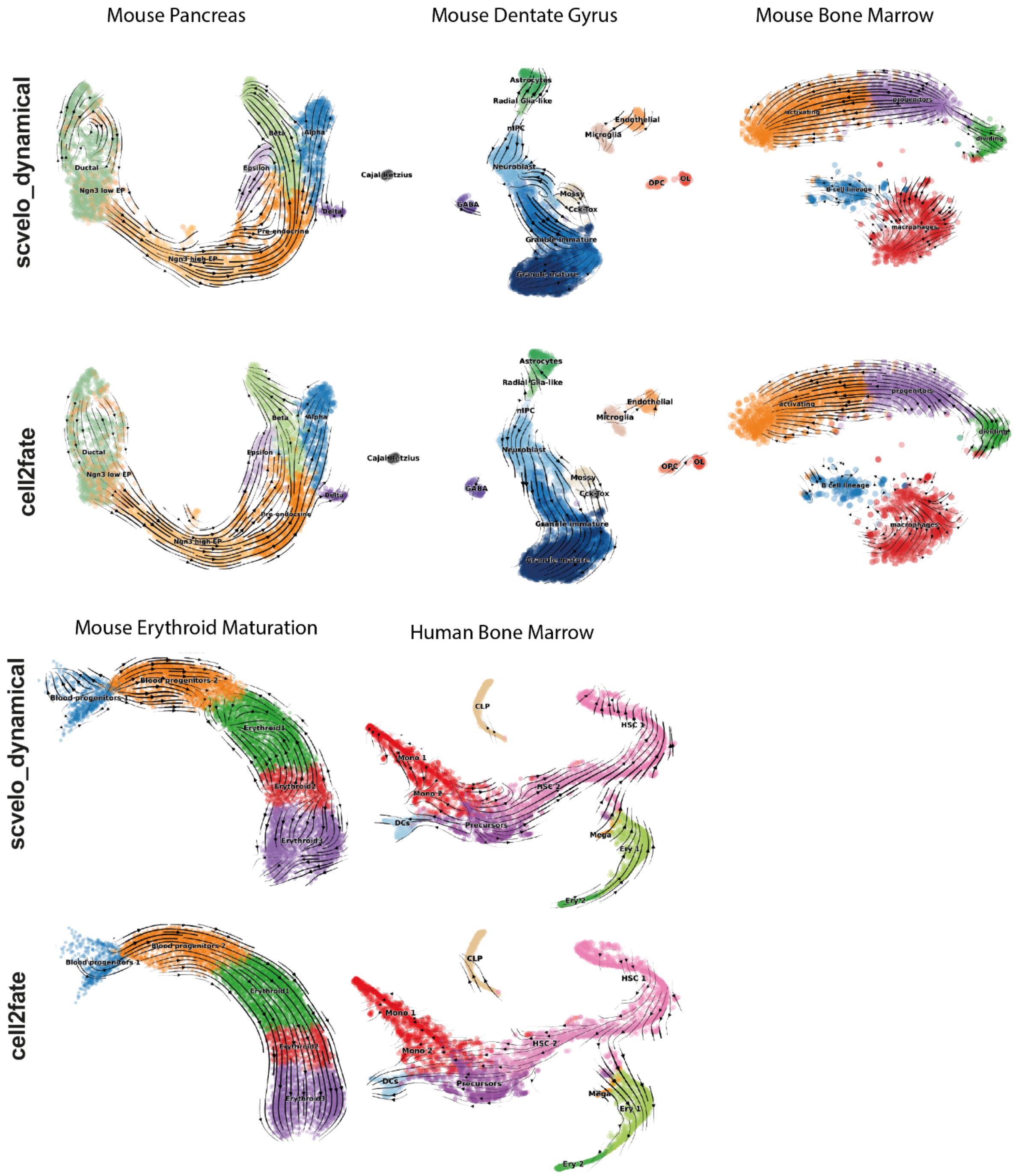


**Supp. Figure 11: all RNA velocity graph UMAP projections for scvelo_dynamical and cell2fate**

*Cell2fate more faithfully captures the expected differentiation trajectories, especially in challenging cases, such as mature granule neurons in the Dentate Gyrus dataset or complex transcription rate changes in the Erythroid Maturation and Human Bone Marrow datasets.*

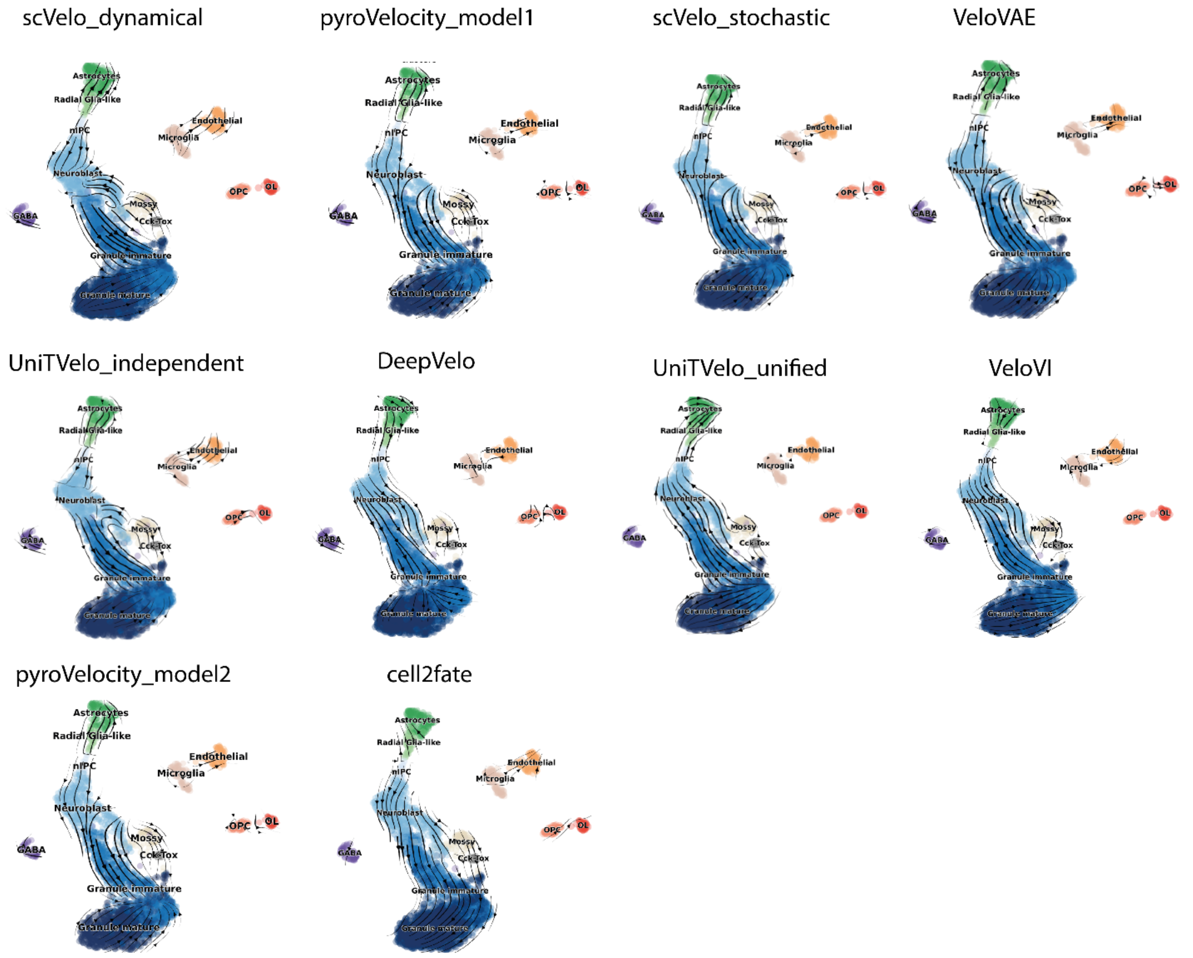


**Supp. Figure 12: velocity graph UMAPs for dentate gyrus data across all benchmarked methods**

*Cell2fate more faithfully captures the expected differentiation trajectories, especially in challenging cases, such as mature granule neurons.*

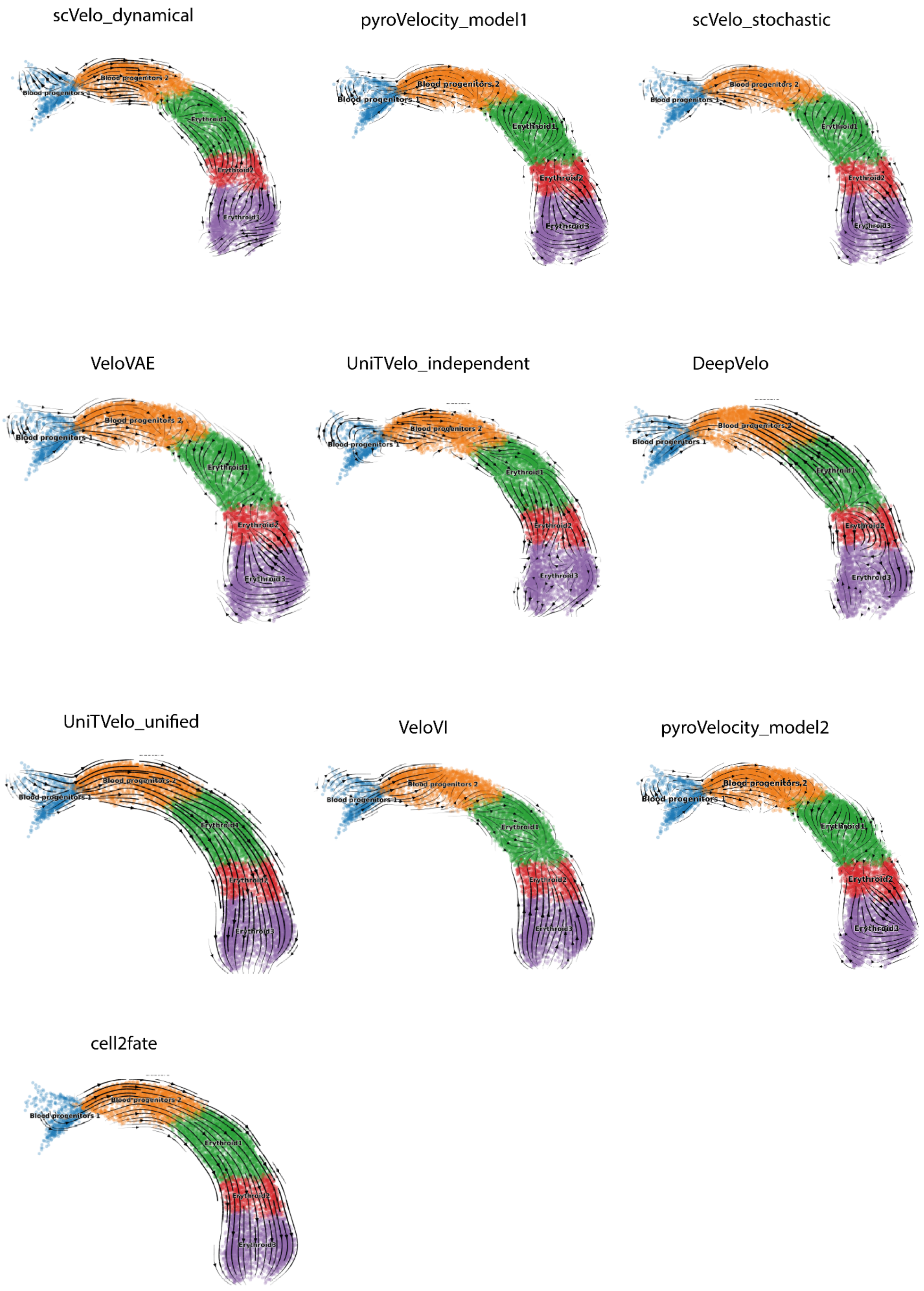


**Supp. Figure 13: velocity graph UMAPs for erythroid maturation data across all benchmarked methods** *UniTVelo and cell2fate produce the expected velocity arrows throughout the trajectory.*

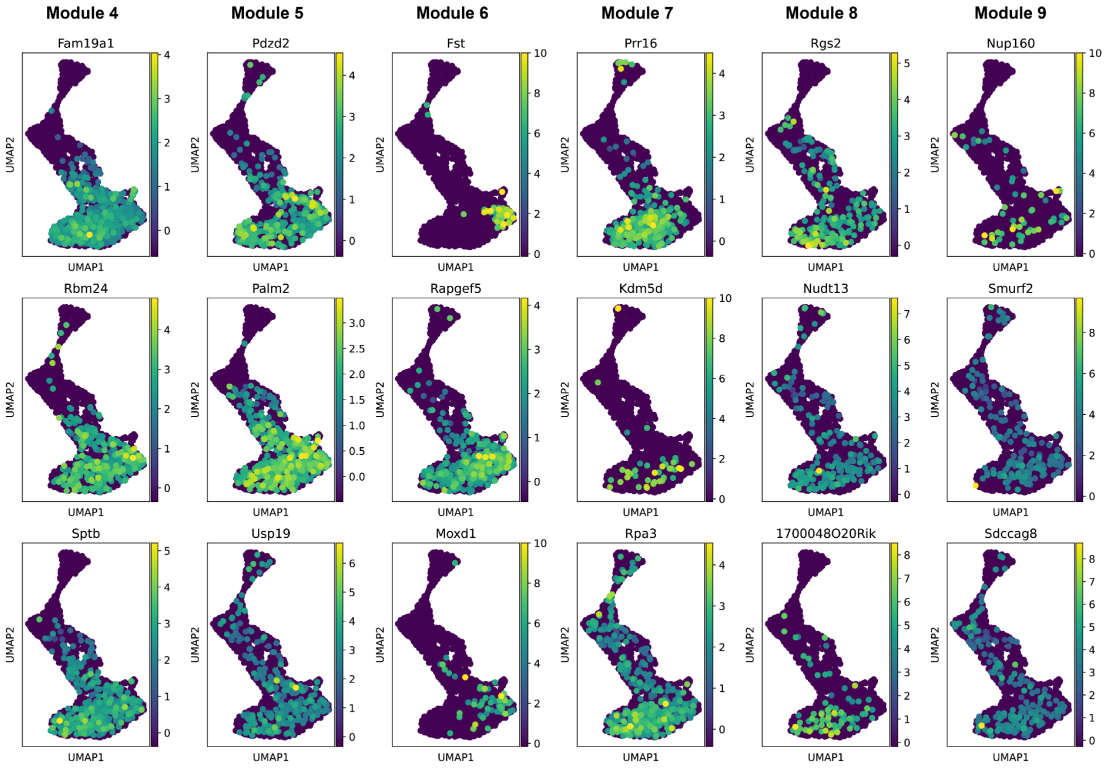


**Supp. Figure 14: Module marker genes in Dentate Gyrus data** *Modules are ordered from left to right, the most significant module gene is at the top, followed by the second and third*

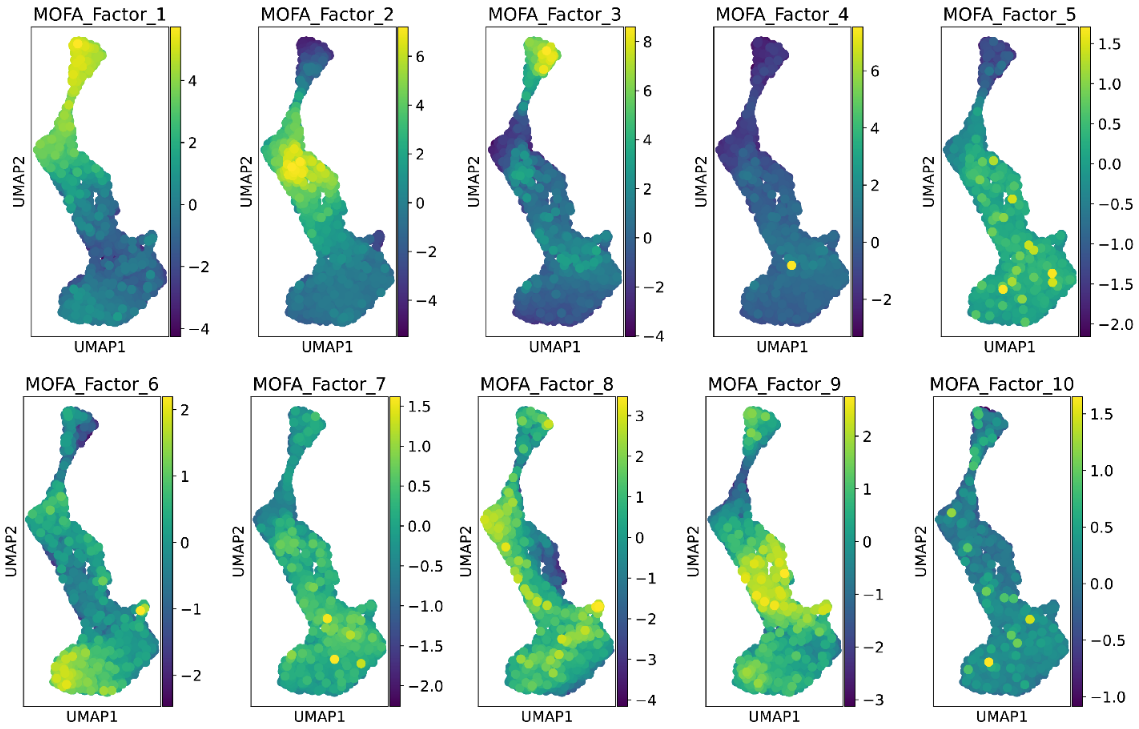


**Supp. Figure 15: Dentate Gyrus data factors obtained with MOFA factors** *Some MOFA factors associate with the differentiation trajectory in this dataset, while others are more broadly distributed.*

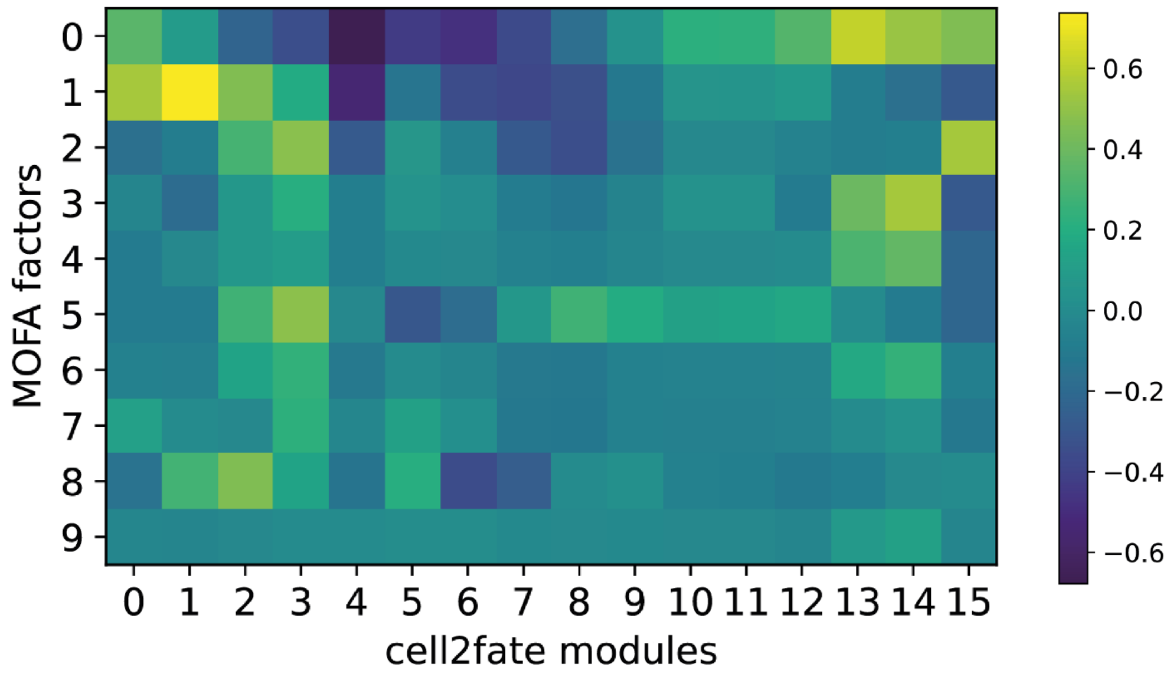


**Supp. Figure 16: Correlations between gene loadings of MOFA factors and cell2fate modules** *Correlations are low throughout, with particularly low values for modules 4 to 9 that are active in neurons. This indicates that cell2fate and MOFA find largely distinct expression dimensions.*

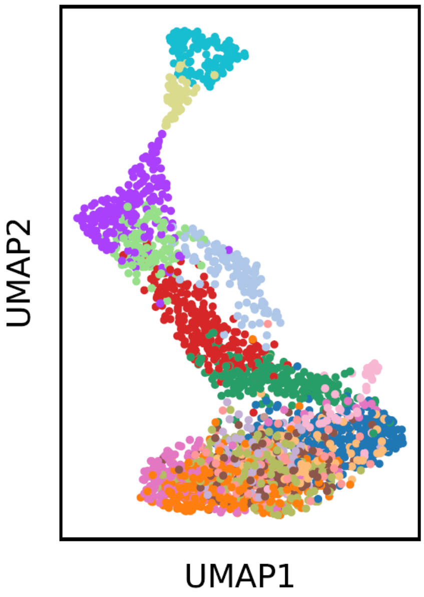


**Supp. Figure 17: Leiden clustering of dentate gyrus data with resolution parameter set to 2** *Leiden clusters partition the differentiation trajectory of Astrocytes (top of UMAP) neuronal progenitors and neurons (middle), but are diffusely distributed across mature granule neurons (bottom).*

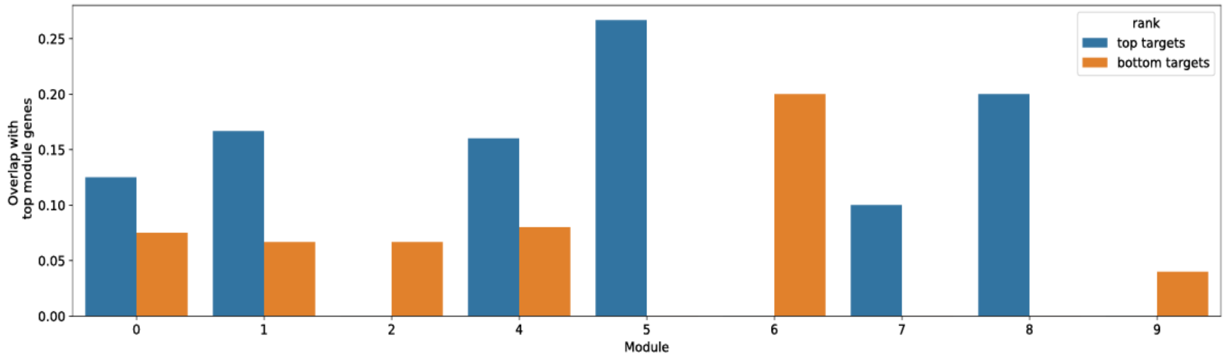


**Supp. Figure 18: overlap of top 10 and bottom 10 binding targets of top 20 module transcription factors with top 300 module genes.** *Putative promoter sequences were first extracted in the genomic vicinity of the top 300 module genes of each module. Binding affinities were then predicted between these sequences and top 20 module transcription factors using the ProBound*[25] *algorithm.*

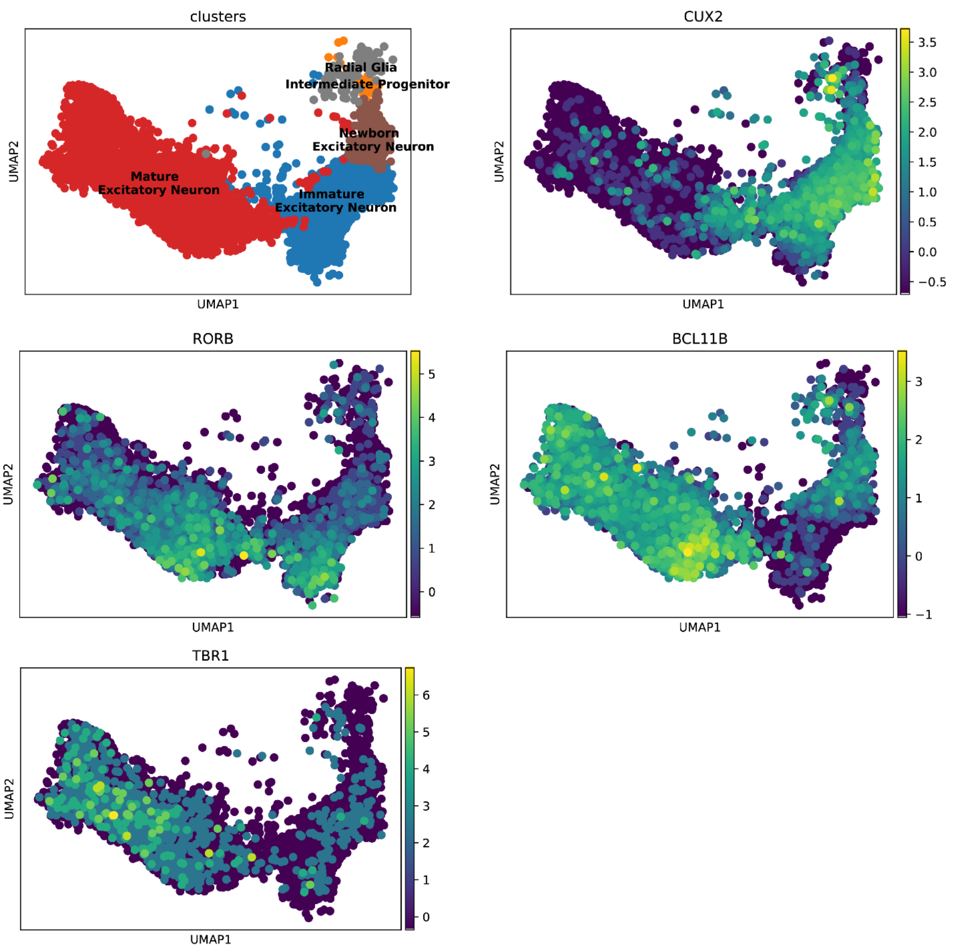


**Supp. Figure 19: neuronal subtype markers visualised in human developing brain data UMAP**

*(Correspondence between subtypes and markers is as follows: CUX2 = upper layer, RORB = layer 4, BCL11B = layer 5, TBR1 = layer 6)*

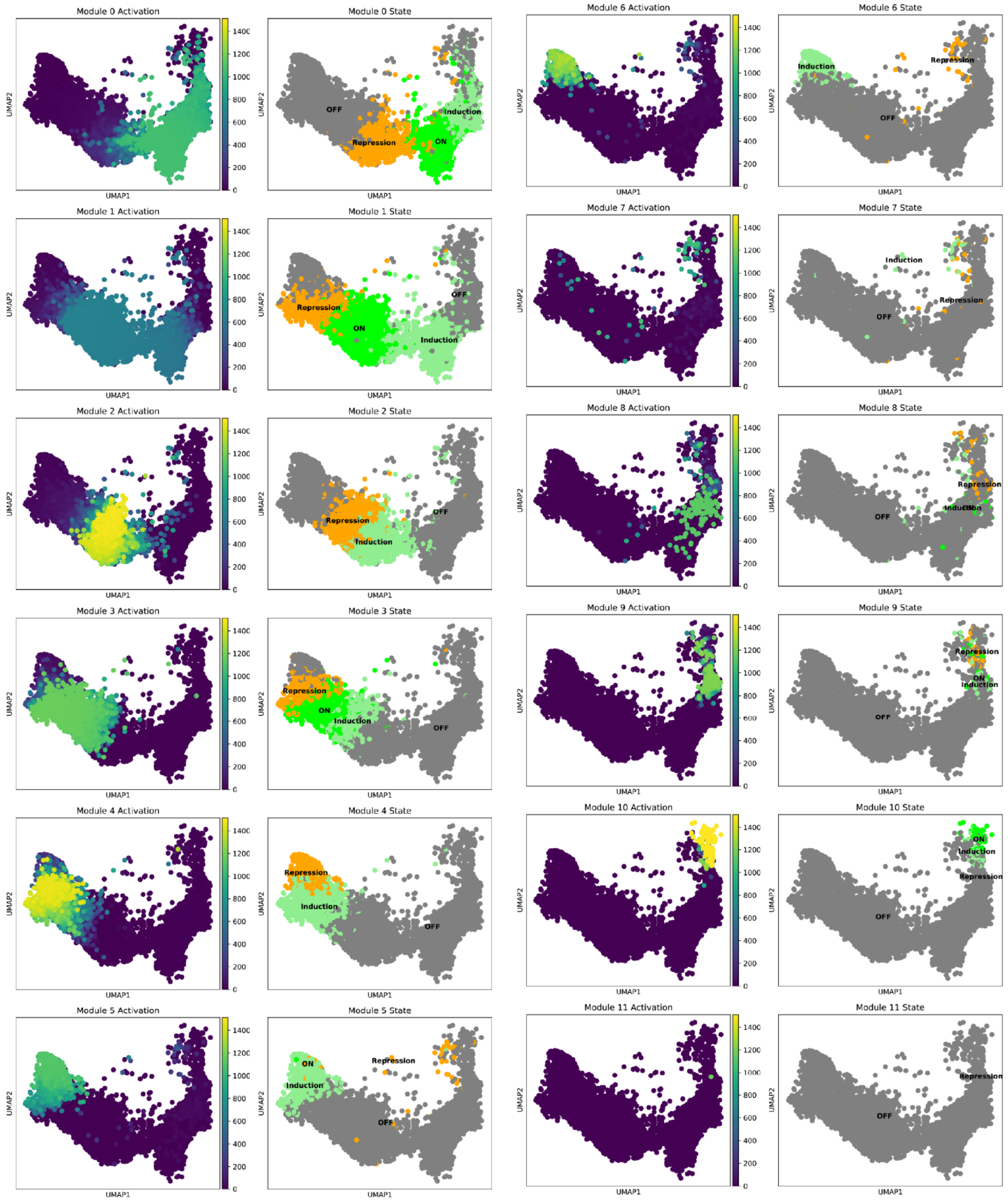


**Supp. Figure 20: module activation and state UMAP plots for human brain data**

*UMAPs coloured by module activation in columns 1 and 3 and state in columns 2 and 4, highlight a sequence of transcriptional programs switched on during differentiation.*

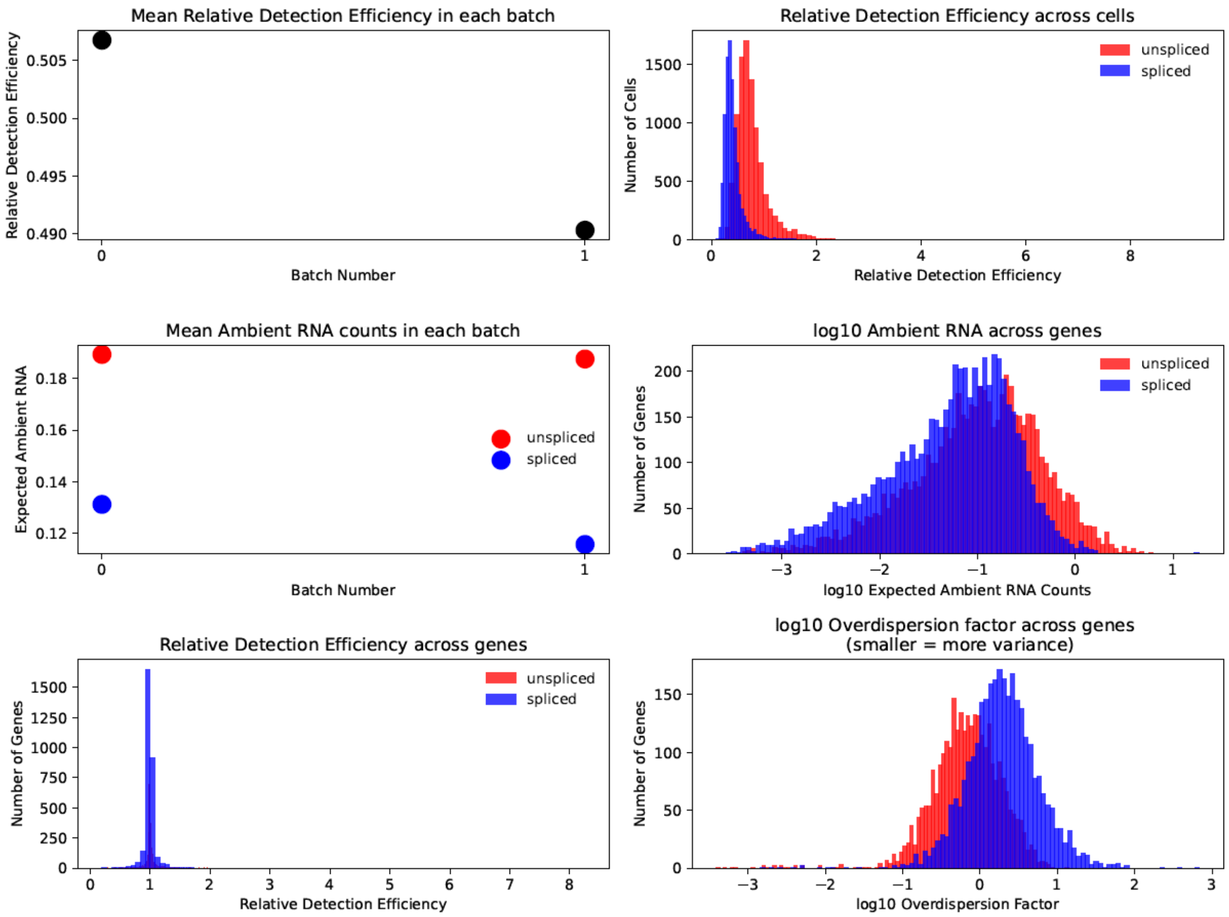


**Supp. Figure 21: overview of noise and technical variables for human brain data**

*In contrast to cell2fate model fits on single-cell data, unspliced counts are predicted to have higher detection efficiency than spliced counts in this single-nucleus dataset (top right). However, they still have higher ambient RNA (middle right) and more noise, corresponding to a lower overdispersion parameter (bottom right).*

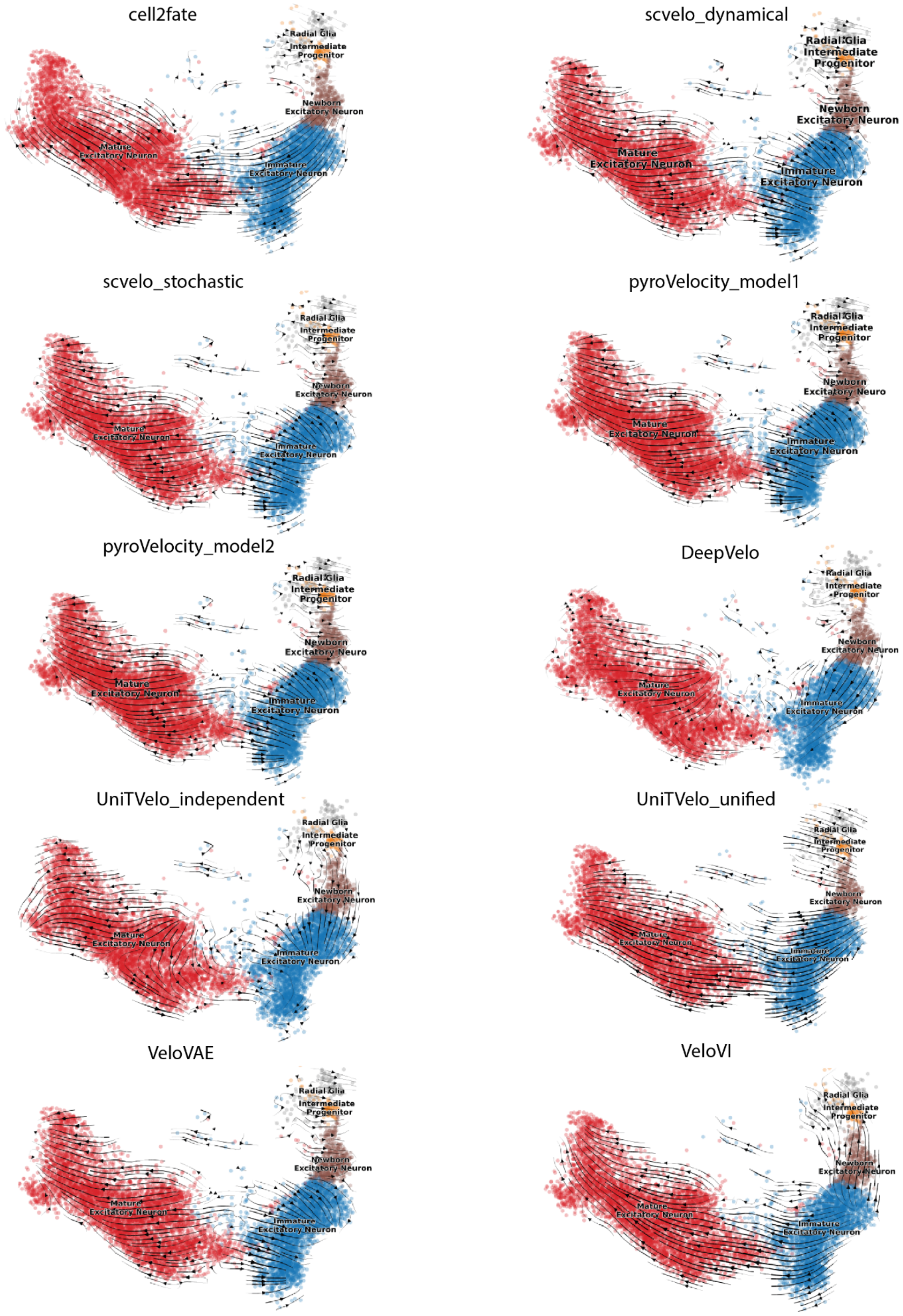


**Supp. Figure 22: Velocity Graph UMAP projections for all 10 RNA velocity methods for human developing brain data**

*cell2fate accurately captures the differentiation trajectory from progenitors (right) to mature neurons (left).*

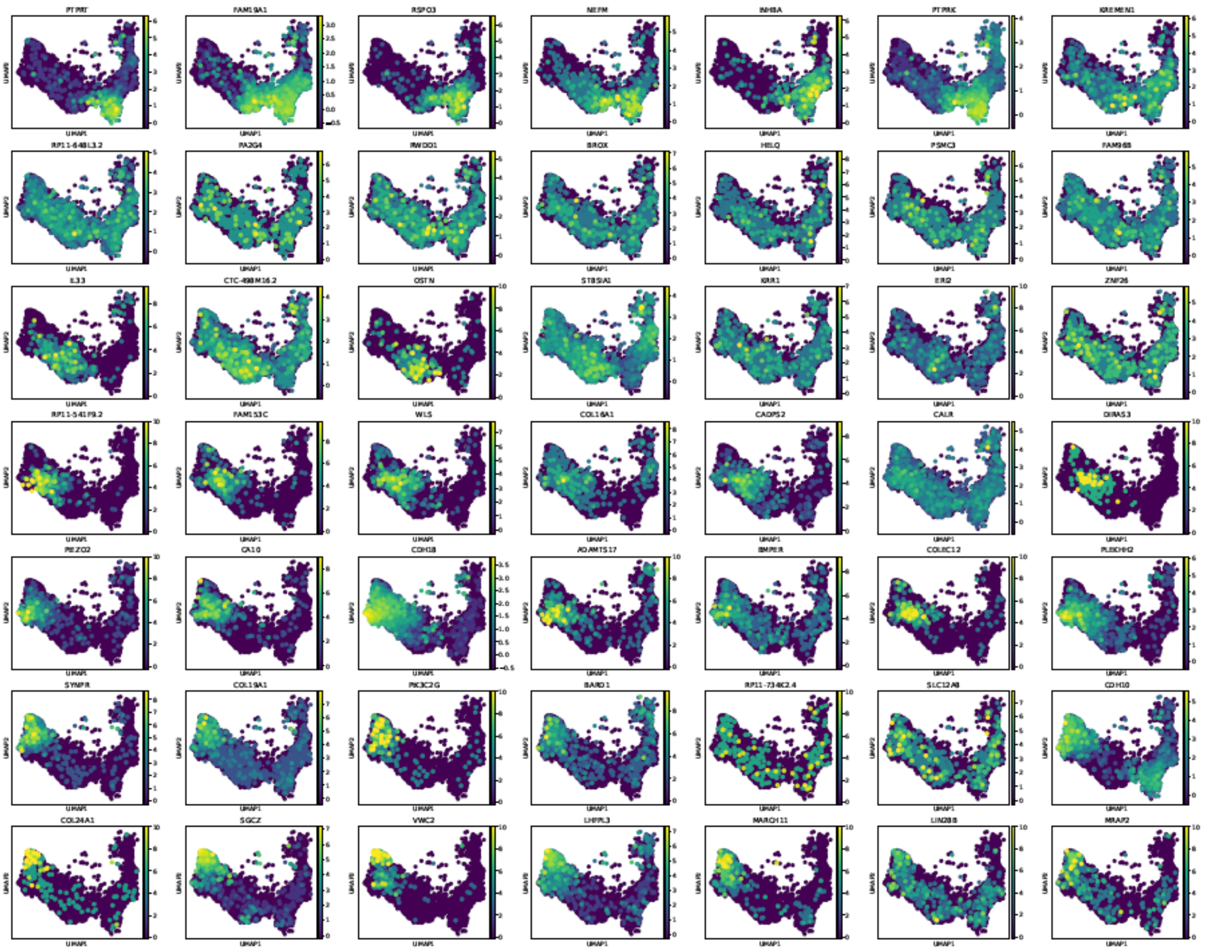


**Supp. Figure 23: top 7 marker genes (left to right) for modules 0 (top) to 6 (bottom) in human developing brain data**

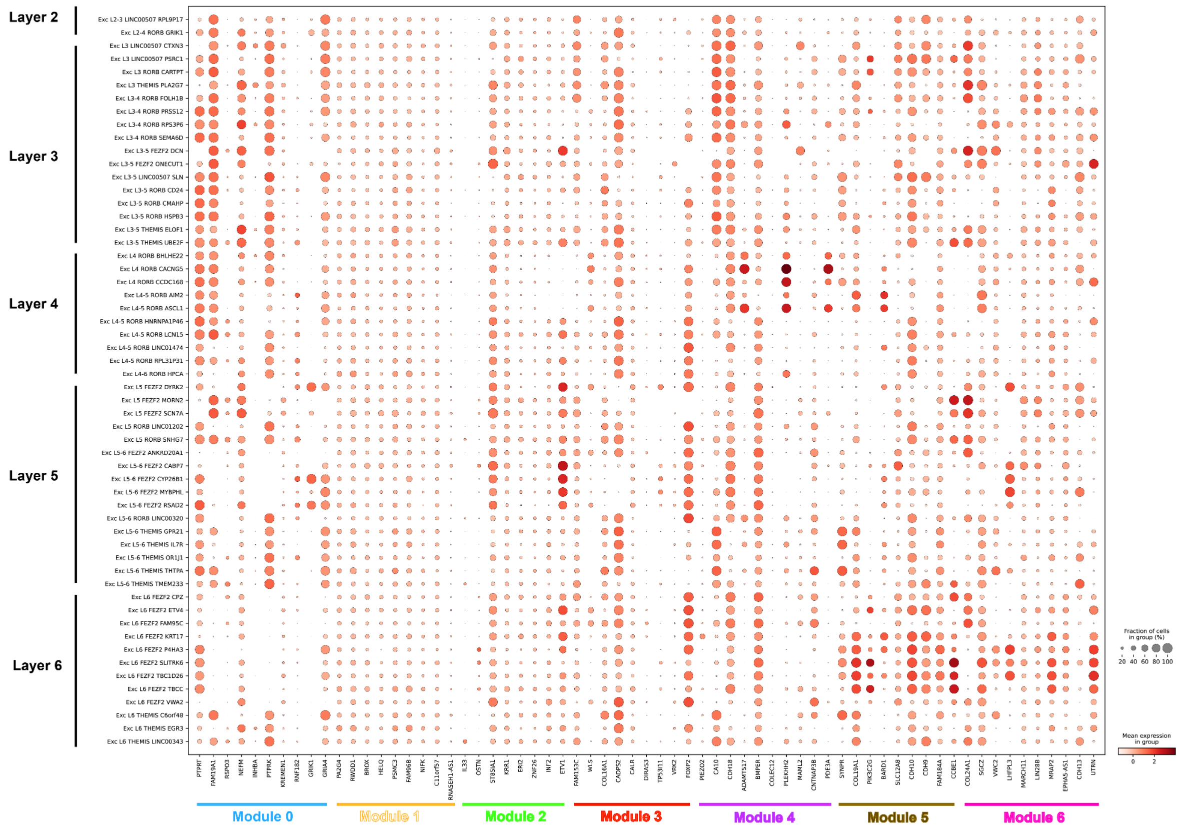


**Supp. Figure 24: Expression of top 10 markers of all 7 neuron differentiation modules in adult human brain data[28]**

*Cell2fate module markers (left to right) are mostly expressed across all neuronal subtypes (top to bottom), indicating that cell2fate modules mostly capture a pan-neuronal differentiation trajectory.*

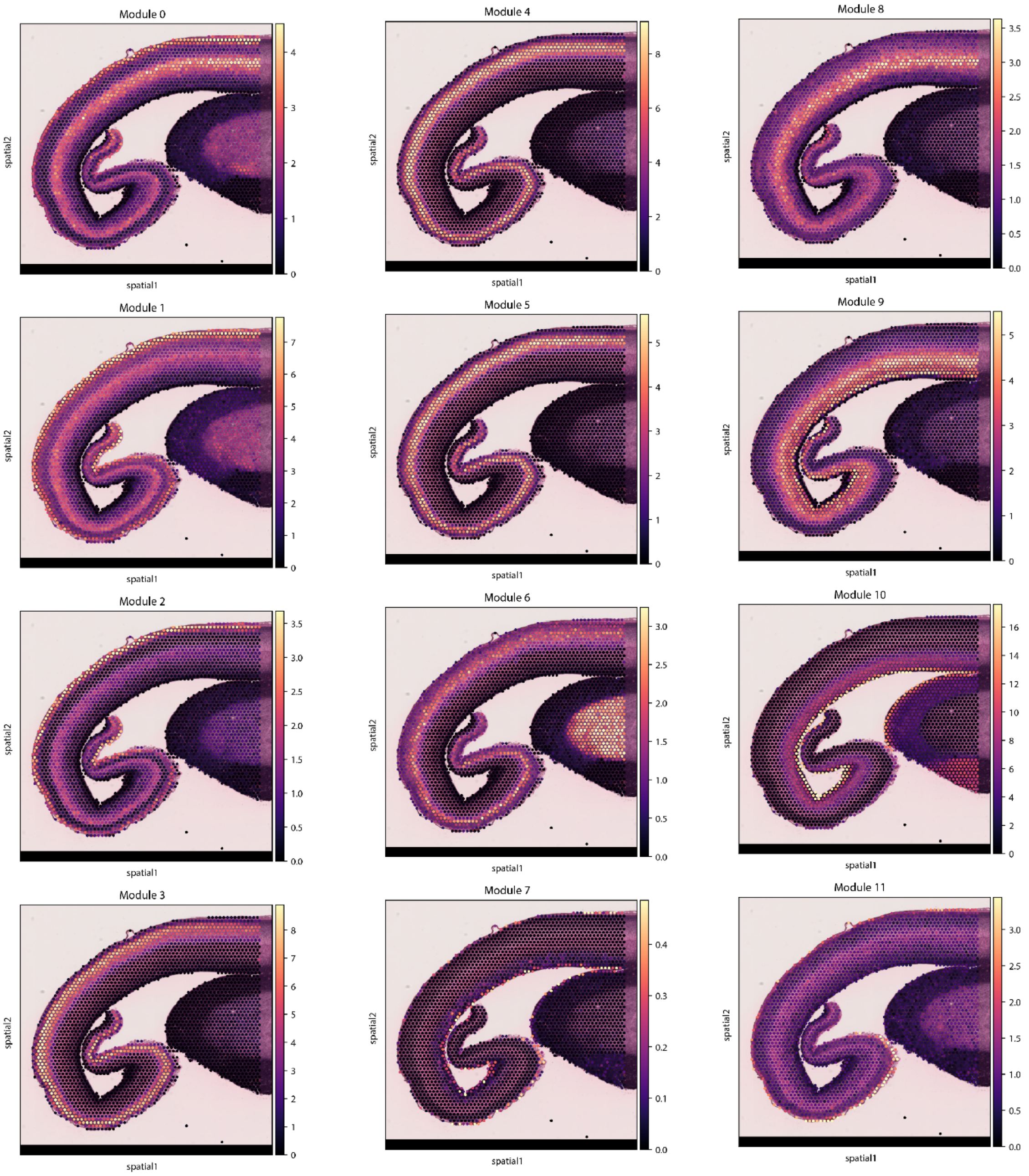


**Supp. Figure 25: cell2location mapping for all cell2fate modules in human developing brain data**

*Cell2fate modules are mapped to distinct locations. Immature neuron modules (e.g. 0,1,2) map towards the inside of the developing human brain section or into the upper layers, while modules corresponding to more mature neurons (e.g. 4,5) map to deep layers.*
